# Pericytes regulate vascular immune homeostasis in the CNS

**DOI:** 10.1101/644120

**Authors:** Orsolya Török, Bettina Schreiner, Hsing-Chuan Tsai, Sebastian Utz, Johanna Schaffenrath, Sina Nassiri, Mauro Delorenzi, Adriano Aguzzi, May H. Han, Melanie Greter, Burkhard Becher, Annika Keller

## Abstract

Brain endothelium possesses several organ-specific features collectively known as the blood-brain barrier (BBB). In addition, trafficking of immune cells in the healthy central nervous system (CNS) is tightly regulated by CNS vasculature. In CNS autoimmune diseases such as multiple sclerosis (MS), these homeostatic mechanisms are overcome by autoreactive lymphocyte entry into the CNS causing inflammatory demyelinating immunopathology. Previous studies have shown that pericytes regulate the development of organ-specific characteristics of brain vasculature such as the BBB and astrocytic end-feet. Whether pericytes are involved in the control of leukocyte trafficking remains elusive. Using adult, pericyte-deficient mice (*Pdgfb^ret/ret^*), we show here that brain vasculature devoid of pericytes shows increased expression of VCAM-1 and ICAM-1, which is accompanied by increased leukocyte infiltration of dendritic cells, monocytes and T cells into the brain, but not spinal cord parenchyma. Regional differences enabling leukocyte trafficking into the brain as opposed to the spinal cord inversely correlate with the pericyte coverage of blood vessels. Upon induction of experimental autoimmune encephalitomyelitis (EAE), pericyte-deficient mice succumb to severe neurological impairment. Treatment with first line MS therapy - fingolimod significantly reverses EAE, indicating that the observed phenotype is due to the massive influx of immune cells into the brain. Furthermore, pericyte-deficiency in mice that express myelin oligodendrocyte glycoprotein peptide (MOG_35-55_) specific T cell receptor (*Pdgfb^ret/ret^; 2D2^Tg^*) leads to the development of spontaneous neurological symptoms paralleled by massive influx of leukocytes into the brain, suggesting altered brain vascular immune quiescence as a prime cause of exaggerated neuroinflammation. Thus, we show that pericytes indirectly restrict immune cell transmigration into the CNS under homeostatic conditions and during autoimmune-driven neuroinflammation by inducing immune quiescence of brain endothelial cells.

## Introduction

CNS vasculature possesses specific features collectively referred to as the BBB, which localizes to endothelial cells. The BBB ensures the delivery of essential nutrients, while preventing the entry of xenobiotics into the brain. In addition, brain endothelial cells restrict the invasion of leukocytes into brain parenchyma, thus contributing to immune privilege of the CNS. BBB function is induced by neural tissue and established by all cell types constituting the neurovascular unit (NVU). Pericytes and mural cells residing on the abluminal side of capillaries and post-capillary venules, regulate several features of the BBB ^1, 2^. Studies on *Pdgfb* and *Pdgfrb* mouse mutants, which exhibit variable pericyte loss, have demonstrated that pericytes negatively regulate endothelial transcytosis which, if not suppressed, leads to increased BBB permeability to plasma proteins ^1, 2^. In addition, pericyte-deficient vessels show abnormal astrocyte end-feet polarization ^1^. Thus, pericytes regulate several characteristics of brain vasculature during development and in the adult organism ^1, 2^. Whether the non-permissive properties of brain vasculature to leukocyte trafficking in the adult organism are regulated by pericytes has not been addressed. Interestingly, increasing evidence points to the role of pericytes in leukocyte extravasation in peripheral organs such as the skin and the striated muscle, and in tumors ^3–5^.

Increased vascular permeability to plasma proteins and immune cells accompanies neurological disorders such as MS, stroke and Alzheimer’s disease (reviewed in ^6, 7^). In MS, a chronic inflammatory and degenerative neurological disorder ^8^, autoreactive lymphocytes infiltrate CNS parenchyma leading to focal inflammatory infiltrates, demyelination, axonal damage and neurodegeneration. This process is accompanied by increased BBB permeability to plasma. However, it is not known whether vascular damage precedes the formation of inflammatory lesions and influences the spatial distribution of demyelinating lesions or whether infiltrating immune cells induce the BBB dysfunction ^9, 10^. Although enormous efforts have been made to understand the pathophysiology of autoimmunity in MS, knowledge regarding the pathological changes of CNS vasculature that permit extravasation of auto-reactive leukocytes is still limited ^11^. This knowledge gap emphasizes the importance of understanding pathological changes at the NVU, which may facilitate entry of autoimmune T-cells as well as the anatomical localization of lesions.

In this study, we investigate how pericytes regulate immune cell trafficking into the CNS during homeostasis and neuroinflammation. We show, using a pericyte deficient mouse line (*Pdgfb^ret/ret^*), that pericytes maintain an anti-inflammatory vascular phenotype and thus prevent leukocyte extravasation into the brain parenchyma. When neuroinflammation is induced, pericyte deficiency promotes an exaggerated phenotype accompanied by massive immune cell infiltration preferentially into the brain, as opposed to the spinal cord. The MS drug fingolimod rescues the exaggerated phenotype indicating that the severe phenotype in *Pdgfb^ret/ret^* mice is caused by immune cell infiltration into the brain. In addition, pericyte-deficient mice harboring naive MOG peptide-specific T-cells (*Pdgfb^ret/ret^; 2D2^Tg^*) develop cerebellar ataxia and massive infiltration of leukocytes into the brain. We show that regional differences in the permissiveness to leukocyte trafficking into the brain as opposed to the spinal cord inversely correlates with the vessel pericyte-coverage suggesting vasculature of the CNS directing the spatial distribution of neuroinflammation.

## Results

### Pericytes control expression of leukocyte adhesion molecules in the brain vasculature

To address the question of whether pericytes regulate immune cell trafficking into the CNS, we used a pericyte-deficient mouse line - *Pdgfb^ret/ret^*, which shows 89% reduction in brain pericyte numbers and 75% reduction in pericyte vessel coverage in adult animals ^1^. Earlier studies have shown that pericyte-deficiency in embryos leads to increased mRNA levels of leukocyte adhesion molecules (LAMs) on endothelial cells^1, 2^. We analyzed published microarray data of adult *Pdgfb^ret/ret^* brain microvasculature and detected a deregulation of several LAMs, including vascular cell adhesion molecule 1 (VCAM-1) (log2=0.43, p=0.007), in adult brain microvasculature abated in pericyte numbers (Supplementary Fig. 1a). To corroborate these findings, we investigated whether strongly reduced pericyte coverage in adult mice leads to changes of LAMs at the protein level. We focused on the expression of VCAM-1 and intercellular adhesion molecule 1 (ICAM-1), which play a major role in the cascade of immune cell transmigration into tissues ^12^. We detected a zonated endothelial expression of both LAMs in control mice (Fig. 1a, b), similar to a published study ^13^. In brains of *Pdgfb^ret/ret^* mice, the zonated expression pattern was lost and paralleled by a conspicuously stronger staining of VCAM-1 and ICAM-1 (Fig. 1a, b). Quantification of VCAM-1 and ICAM-1 vessel surface coverage in the cerebral cortex and in the striatum showed a significant increase of VCAM-1 and ICAM-1 expression in *Pdgfb^ret/ret^* mice compared to controls (Fig. 1c). Thus, pericyte-deficiency results in an increased expression of LAMs on the brain vasculature also in adult mice.

**Figure 1.**
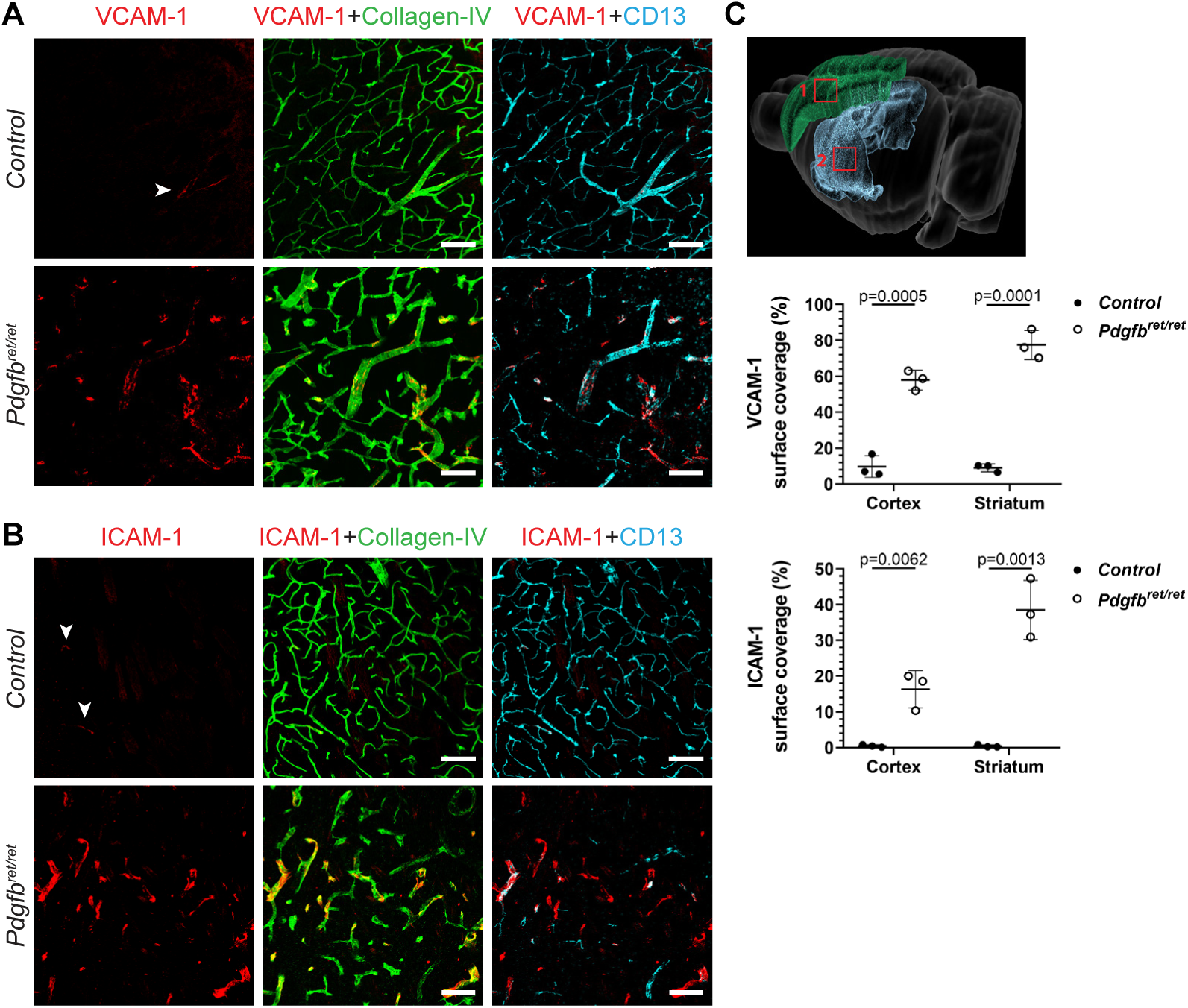
Increased expression of LAMs on pericyte-deficient brain vasculature. Immunofluorescent stainings showing the expression of VCAM-1 (**a**) and ICAM-1 (**b**) on brain vessels in the striatum of control and pericyte-deficient mice (*Pdgfb^ret/ret^*). Vascular basement membrane was visualized with collagen-IV (in green) and pericytes with CD13 (in cyan) immunostaining. Quantification of vascular surface coverage of VCAM-1 and ICAM-1 (**c**) in the cortex and striatum in control and pericyte-deficient mice (*Pdgfb^ret/ret^*). 3D brain image depicting the somatosensory cortex and striatum was generated with the Blue Brain Cell Atlas (bbp.epfl.ch/nexus/cell-atlas/). N=3 mice per genotype. Unpaired t-test was used to determine the statistical significance. Data are presented as the mean ± SD. Scale bars are 100 µm.

### Pericytes control leukocyte extravasation into the brain

We next asked whether increased expression of LAMs on the brain endothelium of pericyte-deficient mice is accompanied by leukocyte infiltration into brain parenchyma. We first analyzed the presence of CD45^hi^ leukocytes in different anatomical regions of the brain by immunofluorescent staining and confocal imaging. Adult *Pdgfb^ret/ret^* mice showed numerous CD45^hi^ leukocyte infiltrates in the brain parenchyma (Fig. 2a). High magnification images revealed that CD45^hi^ cells were found in the vessel lumen, in the brain parenchyma and clustered around blood vessels in the brains of *Pdgfb^ret/ret^* mice, whereas in control mice CD45^hi^ cells were detected but they resided in the lumen of blood vessels (Fig. 2b). Although all cerebral regions in *Pdgfb^ret/ret^* mice showed altered vascular expression of LAMs and immune cell infiltrates, quantification of extravasated leukocytes in different brain regions showed that corpus callosum contained more transmigrated CD45^hi^ cells compared to the striatum and cortex (Fig. 2c). Notably, immunohistochemical analysis of CD45^hi^ cells in the spinal cord of *Pdgfb^ret/ret^* mice showed the absence of immune cell infiltrates in the spinal cord parenchyma (Supplementary Fig. 1b).

**Figure 2.**
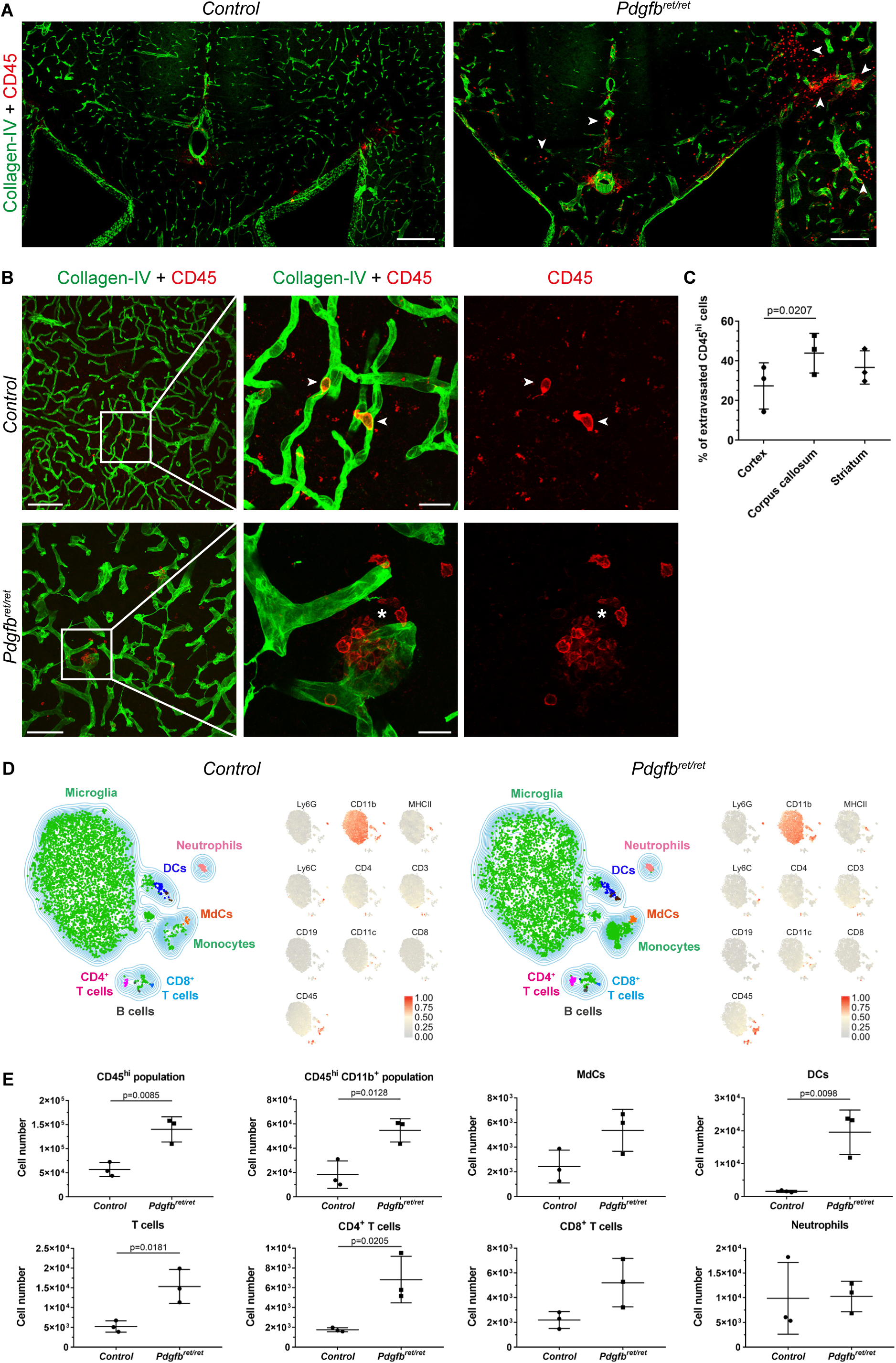
Increased extravasation of leukocytes in the brain parenchyma of pericyte-deficient brain. (**a**) Overview images of the periventricular areas in the brains of control and pericyte-deficient mice (*Pdgfb^ret/ret^*). Arrowheads indicate leukocyte infiltrates (CD45, in red) in the parenchyma of *Pdgfb^ret/ret^* mice. Blood vessels are visualized by collagen IV staining (in green). (**b**) High magnification images showing a parenchymal infiltrate of CD45^hi^ leukocytes (in red, asterix) in the cortex of *Pdgfb^ret/ret^* mice. In control mice, few CD45^hi^ leukocytes (arrowheads) are found only in the lumen of blood vessels (collagen IV, in green). Of note, immunohistochemical sections of control mice were imaged using a higher detector gain (voltage) than images of *Pdgfb^ret/ret^* mice in order to visualize few leukocytes in the lumen of blood vessels. In the control images, in addition to intravascular leukocytes, microglia are detected (CD45^low^). (**c**) Quantification of extravasated CD45^hi^ leukocytes in different anatomical regions in the brains of *Pdgfb^ret/ret^* mice (n=3). Ordinary one-way ANOVA test was used to determine the statistical significance. (**d**) T-SNE plot displaying immune cell populations in the brain and spinal cord of naïve control and mice *Pdgfb^ret/ret^* mice (n=3 per genotype) analyzed by flow cytometry. Data were transformed and percentiles normalized. Median expression values (0-1) of selected markers are shown. (**e**) Quantification of the absolute cell numbers of detected immune cell populations using flow cytometry (n=3 mice per genotype). Unpaired t-test was used to determine statistical significance. Data are presented as the mean ± SD. Scale bars: a - 250 µm, b - 100 µm and magnified insets - 20 µm.

Having established that brain parenchyma of *Pdgfb^ret/ret^* mice contains CD45^hi^ cells, we used flow cytometry to identify immune cell populations. To give an overview of all immune cell populations in the CNS, leukocytes were isolated from the CNS, analyzed by flow cytometry, categorized by unsupervised meta clustering and visualized in a t-distributed stochastic neighbor embedding (t-SNE) map. This approach confirmed the immunohistochemistry findings and showed increased frequencies of CD45^hi^ cells in the CNS of pericyte-deficient mice. The majority of cells increased in *Pdgfb^ret/ret^* mice were CD45^hi^CD11b^+^ myeloid cells (mainly composed of monocytes and CD11c^+^ dendritic cells (DCs)), and T cells (Fig. 2d). Quantification of the absolute cell numbers of manually gated immune cells subsets showed a significant increase in the number of CD45^hi^CD11b^+^ myeloid cells, DCs (3.5% of the CD45^hi^CD11b^+^ population), and CD4^+^ T cell populations compared to controls (Fig. 2e). In addition, we observed an increased number of Ly6C^hi^MHC-II^+^ monocyte derived cells (MdCs) and the CD8^+^ T cell population in the CNS of pericyte-deficient animals compared to controls (Fig. 2e). Subsequent flow cytometry analysis of brain and spinal cord separately confirmed the presence of CD45^hi^ cells (DCs, MdCs, T cells) in brain and the absence of CD45^hi^ cells in the spinal cord of *Pdgfb^ret/ret^* mice (Supplementary Fig. 1c, d).

We next analyzed leukocyte populations in blood as well as in primary and secondary lymphoid organs to ensure that the increased number of leukocytes in the brains of pericyte-deficient animals is not caused by peripheral alterations. The total cell number in thymus, spleen, axillary and inguinal lymph nodes and blood was determined using an automated cell counter with isolated cells stained for further flow cytometry analysis. The cell number in the thymus, spleen, lymph node and blood was comparable between *Pdgfb^ret/ret^* and control mice (Supplementary Fig. 2a). Subsequent analysis of leukocyte populations did not show a skewing between *Pdgfb^ret/ret^* and control mice in blood, lymph nodes and spleen (Supplementary Fig. 2b, c, d). However, the spleens of *Pdgfb^ret/ret^* mice showed slightly elevated numbers of CD8^+^ T cells compared to controls (Supplementary Fig. 2c). There was no difference in the total leukocyte or in the neutrophil count in blood (Supplementary Fig. 1b), indicating the absence of systemic inflammation in *Pdgfb^ret/ret^* mice. Histological examination of lymphoid organs did not show any differences in the spatial organization of T and B cells (Supplementary Fig. 2f-h) between *Pdgfb^ret/ret^* and control mice. Thus, the increased number of infiltrated leukocyte subsets in the brain of pericyte-deficient mice is not due to increased numbers in the blood or lymphoid organs.

Taken together, our data show that in the absence of pericytes, the adult brain vasculature becomes permissive for leukocyte entry and that the infiltrated leukocyte population consists mostly of dendritic cells, MdCs and T cells.

### Spatial differences in pericyte coverage in the CNS of *Pdgfb^ret/ret^* mice

Previous studies have shown a negative correlation between pericyte coverage and BBB permeability in the brain ^1, 2, 14–16^. We therefore asked whether selective leukocyte infiltration into the brain in *Pdgfb^ret/ret^* (Supplementary Fig. 1b-d) mice can be explained by differences in capillary pericyte coverage in the brain and spinal cord. Immunofluorescent staining of vasculature and pericytes revealed a reduced pericyte coverage on blood vessels in the spinal cord of *Pdgfb^ret/ret^* mice compared to controls. However, capillary pericyte coverage in the spinal cord in *Pdgfb^ret/ret^* mice was more complete when compared to different brain regions (cortex and striatum) (Fig. 3a). In sharp contrast to the brain vasculature, the pattern and morphology of spinal cord vasculature appeared similar to control mice (Fig. 3a). Quantification of vessel surface pericyte coverage in the spinal cord showed that *Pdgfb^ret/ret^* mice have a significantly reduced capillary pericyte coverage compared to control animals (Fig. 3b). However, the observed ~26% reduction of pericyte coverage in spinal cord vasculature of *Pdgfb^ret/ret^* mice is notably less than the previously reported reduction of pericyte coverage in the cortex or deep brain regions (~75 %)^7, 1, 14^. Finally, we investigated whether higher capillary pericyte coverage in the spinal cord of *Pdgfb^ret/ret^* mice parallels normalized expression of VCAM-1 and ICAM-1. Indeed, the expression of VCAM-1 and ICAM-1 on spinal cord vasculature showed a similar zonal expression pattern in control and *Pdgfb^ret/ret^* mice (Fig. 3c, d). Based on these data, we conclude that regional differences in the degree of capillary pericyte coverage in the CNS determine the extent to which brain vasculature expresses LAMs and thereby leukocyte entry into the CNS.

**Figure 3.**
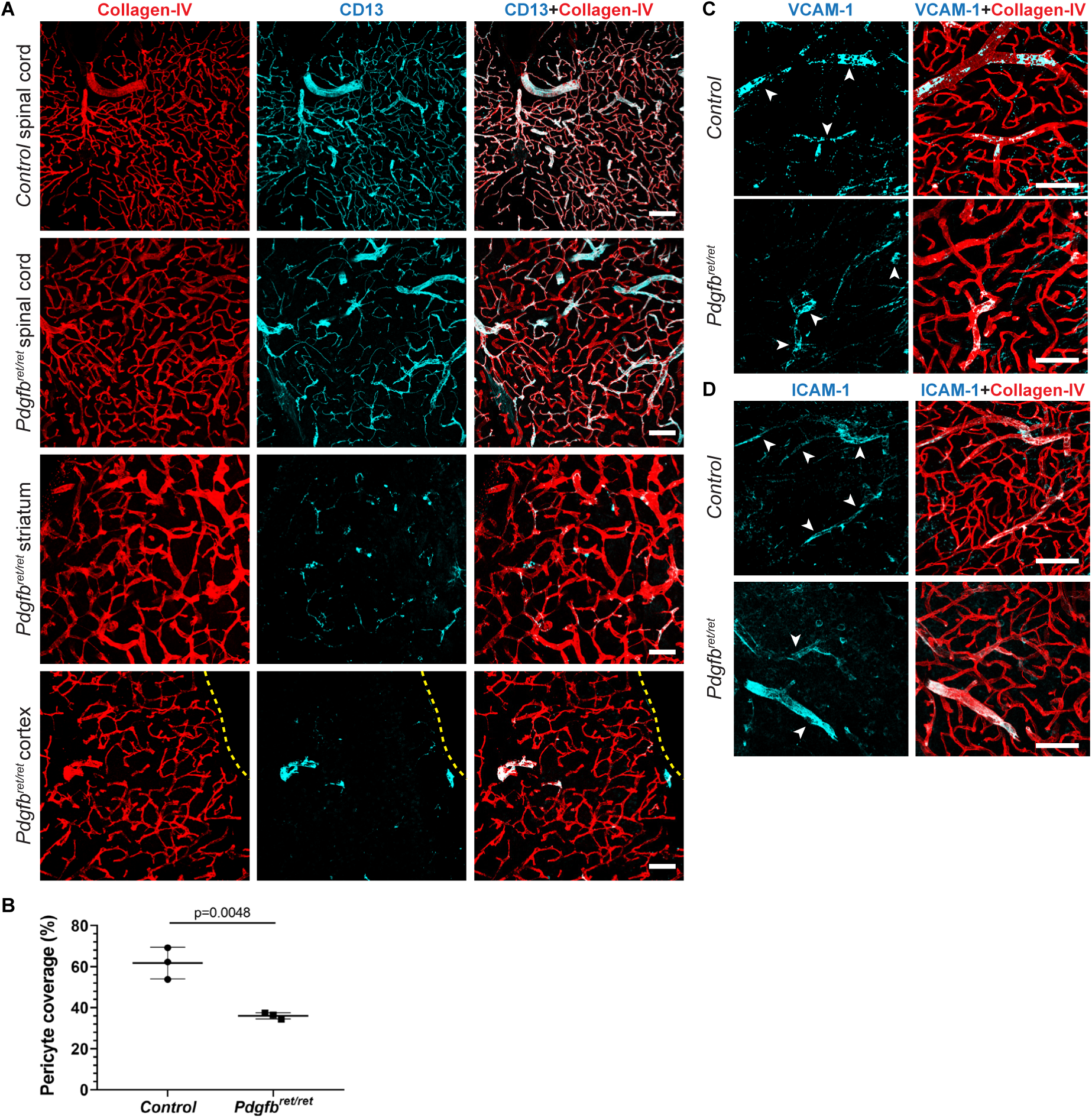
Increased pericyte-coverage and reduced expression of LAMs in the spinal cord vessels of *Pdgfb^ret/ret^* mice. (**a**) Immunofluorescent staining of pericytes (CD13, in cyan) and vasculature (collagen-IV, in red) in different anatomical regions of the CNS (spinal cord, striatum, cortex) in control and *Pdgfb^ret/ret^* mice. The yellow dotted line outlines the cortical surface. (**b**) Quantification of vessel pericyte coverage in the spinal cord of control and *Pdgfb^ret/ret^* mice. N= 3 mice per genotype. VCAM-1 (**c**) and ICAM-1 (**d**) expression (in cyan) on the blood vessels (in red, collagen-IV) in the spinal cord of control and *Pdgfb^ret/ret^* mice. Scale bars are 100 µm. Unpaired t-test was used to determine statistical significance.

### Loss of pericytes does not alter myelin integrity

We next asked whether increased BBB permeability to plasma proteins ^1, 2^ and leukocytes leads to subclinical demyelination in the brains of pericyte-deficient mice. Myelin was visualized luxol fast blue – periodic acid Schiff (LFB-PAS) histochemical stains as well as immunofluorescent labelling with anti-myelin basic protein (MBP) antibody (Supplementary Fig. 3a, c). Quantification of LFB-PAS and anti-MBP staining intensity in the region of high myelin content, the corpus callosum, did not differ between control and *Pdgfb^ret/ret^* mice (Supplementary Fig. 3b, d). A few brain sections of *Pdgfb^ret/ret^* mice stained with LFB-PAS had a reduced staining intensity (score1) in the corpus callosum due to accompanying brain edema ^1^; however, demyelinating lesions were absent. Additionally, we did not detect differences in the ultrastructure of the myelin sheath between *Pdgfb^ret/ret^* and control mice (Supplementary Fig. 3e). Thus, infiltrated leukocytes in brain parenchyma in pericyte-deficient mice do not initiate demyelinating pathology.

### Pericyte-deficient mice present with an aggravated, atypical EAE phenotype

We next investigated whether leukocyte permissive vasculature modifies the course of autoimmune neuroinflammation. In order to address this question, we induced EAE, an animal model of MS ^17^ in control and *Pdgfb^ret/ret^* mice. After active induction of EAE, which replicates both the induction and effector phase of the disease, *Pdgfb^ret/ret^* mice presented with a severe, early onset (4-5 day p.i.) atypical phenotype as well as reduced survival (Fig. 4a, b). We confirmed that control animals in the study (*Pdgfb^wt/ret^*) did not differ from wild-type littermates in the clinical course of EAE (Supplementary Fig. 4a, b). Therefore, *Pdgfb^wt/ret^* mice continued to be used as controls. All control mice presented typical spinal cord EAE signs with ascending paralysis starting at the distal tail. All *Pdgfb^ret/ret^* mice invariably developed the atypical phenotype, which consisted of prominent cerebellar ataxia and spasticity, without ascending paralysis. To score the clinical severity of EAE in *Pdgfb^ret/ret^* mice (Fig. 4b), we adopted an ataxia scoring protocol described by Guyenet et al. ^18^. We noticed that *Pdgfb^ret/ret^* mice presented with a basal ataxia score of 2 already at day 0, which consisted of kyphosis (score 1) and hindlimb clasping (score 1) (Fig. 4b, see “Material and Methods” for the scoring protocol). Adoptive transfer (passive) EAE resulted in the same aggravated atypical EAE in *Pdgfb^ret/ret^* mice as seen in active EAE (Fig. 4c), indicating that the severe phenotype is not due to a pathologically enhanced induction phase in pericyte-deficient mice. Immunization with a non-CNS antigen (ovalbumin peptide), using the same adjuvant, did not result in clinical deficits neither in control nor in *Pdgfb^ret/ret^* mice (data not shown).

**Figure 4.**
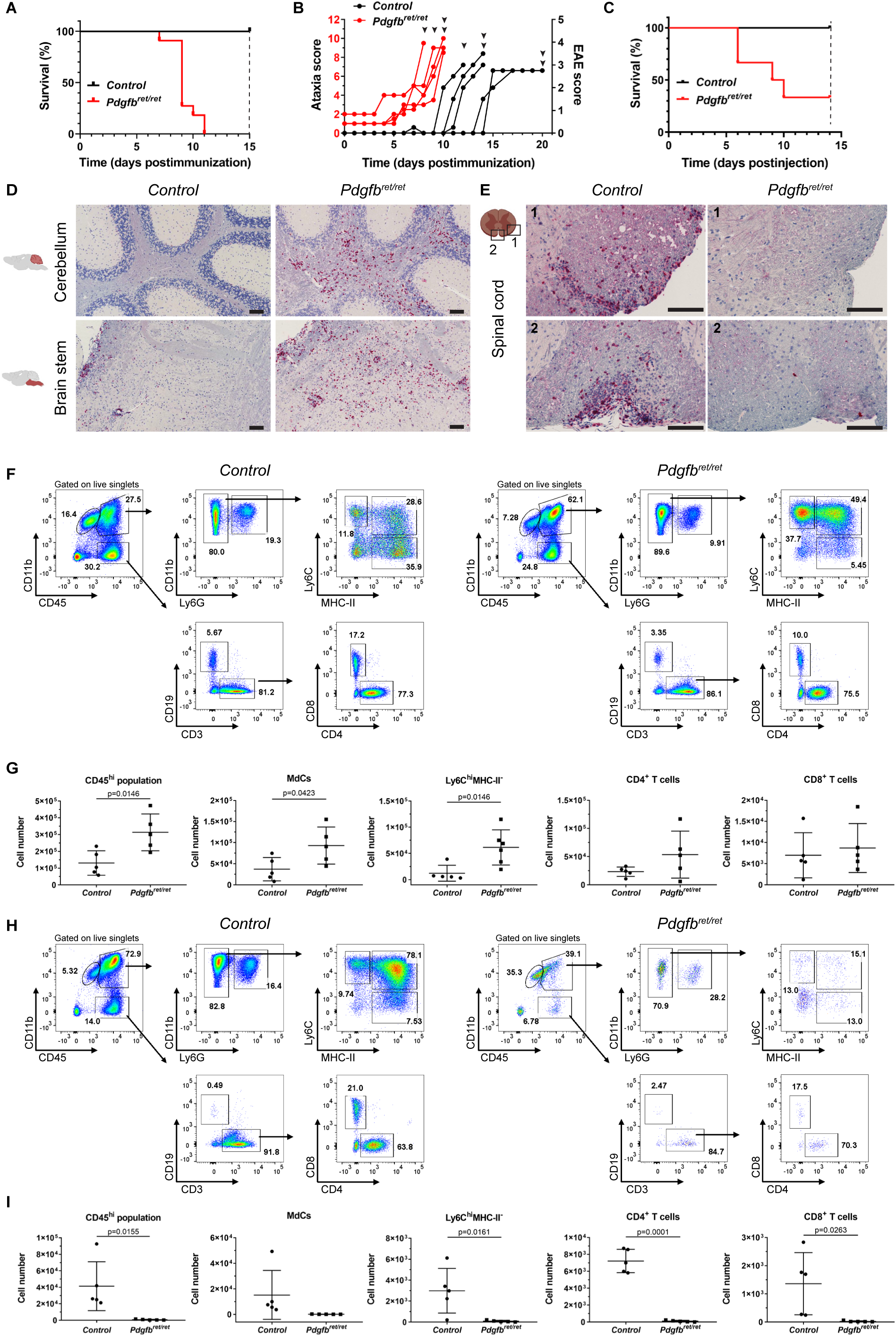
Pericyte-deficient mice succumb to atypical EAE accompanied by increased leukocyte infiltration into the brain. (**a**) Kaplan-Meier survival curves after active induction of EAE. The experiment was terminated on day 15, indicated by black dashed line. Pooled data from two individual experiments. N=11 mice per genotype. Survival curves showed statistical difference (p<0.0001, log-rank test). (**b**) Scoring of neurological symptoms during the course of active EAE. The left y axis shows cerebellar ataxia scores of *Pdgfb^ret/ret^* mice (in red) and the right y axis,classical EAE scores of control mice (in black). Arrowheads indicate when individual mice were sacrificed for flow cytometry analysis. See Materials and Methods for detailed termination criteria. Each line represents symptoms of an individual mouse. N=5 mice per genotype, showing two pooled experiments. (**c**) Kaplan-Meier survival curves after passive induction of EAE. The experiment was terminated on day 14, indicated by black dashed line. Controls - n=5, *Pdgfb^ret/ret^* - n=6). Survival curves showed a statistically significant difference (p=0.0300, log-rank test). (**d**) Immunohistochemical staining of T cells (CD3, in red) of sagittal brain sections of the cerebellum and brain stem after active induction of EAE of control (on day 16 postimmunization) and *Pdgfb^ret/ret^* mice (on day 11 postimmunization). (**e**) Immunohistochemical staining of T cells (CD3, in red) on coronal sections of the spinal cords showing two regions (1, 2) after active induction of EAE in control (on day 16 postimmunization) and *Pdgfb^ret/ret^* mice (on day 11 postimmunization). Tissue sections were counterstained with hematoxylin (**d**, **e**). Representative flow cytometry pseudocolor plots showing the manual gating of microglia and other immune cell populations in the brain (**f**) and spinal cord (**g**) after active induction of EAE of control (EAE score 3) and *Pdgfb^ret/ret^* (ataxia score 9) mice. Quantification of the absolute cell numbers of different leukocyte populations (gated as shown in f and g) in the brains (**h**) and spinal cords (**i**) of control (EAE score 3-3.5) and *Pdgfb^ret/ret^* (ataxia score 9-10) mice using flow cytometry. Shown are pooled data from two individual experiments. N= 5 mice per genotype. Data are presented as the mean ± SD. Unpaired t-test was used to determine the statistical significance. Scale bars are: **d** and **e** – 100 µm.

We next investigated the spatial distribution of infiltrating cells in the CNS after induction of EAE using immunohistochemistry. This analysis showed an increased leukocyte infiltration into the brain parenchyma (cerebral cortex, striatum, corpus callosum, cerebellum, brain stem) of *Pdgfb^ret/ret^* animals, whereas immune cell infiltrates were mostly found in the spinal cord in control animals (Fig. 4d, e; Supplementary Fig. 3c, d). The spinal cords of pericyte-deficient animals were devoid of T-cell infiltrates consistent with the atypical clinical phenotype (Fig. 4e) and with relatively complete vessel pericyte coverage (Fig. 3). Assessment of myelin damage after the induction of EAE showed pronounced demyelination in the brains of pericyte-deficient mice compared to control animals (Supplementary Fig. 3e, f).

We next analyzed which immune cells infiltrate the CNS after active immunization in control and *Pdgfb^ret/ret^* mice. Flow cytometry analysis confirmed the immunohistochemistry results showing an increased number of CD45^hi^ leukocytes in the brains in *Pdgfb^ret/ret^* mice compared to controls (Fig. 4f, g). The majority of these infiltrates (approx. 60 % of live singlets) in *Pdgfb^ret/ret^* mice were CD45^hi^CD11b^+^ myeloid cells (Fig. 4f). Within this population, we detected a significant increase of MdC and Ly6C^hi^MHC-II^−^ subpopulations in the brains of *Pdgfb^ret/ret^* mice compared to controls (Fig. 4g). In addition, there was a trend towards an increased number of CD4^+^ T cells in the brains of *Pdgfb^ret/ret^* mice (Fig. 4g), but no difference in total numbers of neutrophils or B cells (Supplementary Fig. 3g). In agreement with the clinical deficits and immunohistochemistry, the spinal cord of *Pdgfb^ret/ret^* mice was essentially devoid of leukocytes compared to controls (Fig. 4h, i; Supplementary Fig. 3h).

Thus, *Pdgfb^ret/ret^* mice develop a severe, atypical EAE phenotype and show spatially restricted infiltration of inflammatory cells predominantly into brain, consisting mostly of MdC and Ly6C^hi^MHC-II^−^ myeloid cell populations.

### MS drug fingolimod (FTY-720) ameliorates the severe atypical EAE phenotype of *Pdgfb^ret/ret^* mice

We next addressed whether the aggravated phenotype of *Pdgfb^ret/ret^* mice after induction of EAE is caused by the massive influx of peripheral immune cells into the CNS. Mice were treated daily, starting on day 4 post-immunization, with FTY-720 (Fingolimod), a functional antagonist of sphingosine-1-phosphate receptor 1 (S1P1), which causes leukopenia by blocking the egress of lymphocytes from lymph nodes ^19^. All vehicle-treated *Pdgfb^ret/ret^* mice reached termination criteria (ataxia score 8.5-10) after EAE induction whereas FTY-720 treated *Pdgfb^ret/ret^* mice did not develop symptoms of EAE during the course of the experiment (25 days) (Fig. 5a, b). Of note, the ataxia score 2 observed in all *Pdgfb^ret/ret^* mice before FTY-720 administration was not alleviated by FTY-720 treatment. As expected, the EAE score of vehicle-treated control animals improved by day 25 postimmunization. In addition, FTY-720 treated control mice did not develop EAE (Fig. 5b). Flow cytometry analysis of peripheral blood on day 12 post-immunization with MOG peptide confirmed the FTY-720 treatment-induced leukopenia in control and *Pdgfb^ret/ret^* mice (Supplementary Fig. 5a-c). The brains and spinal cords of vehicle treated mice were analyzed with FTY-720 treated mice on the same day for the presence of immune cells when they reached termination criteria (ataxia score 8.5-10) using flow cytometry. In parallel, the brain and spinal cords of control mice (EAE score 3-3.5) were analyzed together with FTY-720 treated controls on the same day. As expected, FTY-720 treated animals had significantly lower numbers of CD45^hi^ immune cells in the CNS both in control and *Pdgfb^ret/ret^* mice (Fig. 5c-f). In addition to reduced number of CD4^+^ and CD8^+^ T cells, we also observed a reduction of myeloid cells (MdC and Ly6C^hi^MHC-II^−^ cells) after FTY-720 treatment in brains of *Pdgfb^ret/ret^* mice (Fig. 5c-e). FTY-720 treatment after the induction of EAE has been shown to reduce the number of circulating monocytes in addition to T-cells ^20^, which could explain the significantly reduced myeloid cells in the spinal cord of control mice (Fig. 5d, f) as well as in the brains of both control and *Pdgfb^ret/ret^* mice (Fig. 5c, e). Immunohistochemical analysis confirmed the reduced infiltration of leukocytes into the brain of pericyte-deficient mice after FTY-720 treatment (Supplementary Fig. 5d). Thus, we conclude that the severe clinical phenotype of pericyte-deficient mice after induction of EAE is caused by excessive entry of peripheral immune cells into the brain and neuroinflammation.

**Figure 5.**
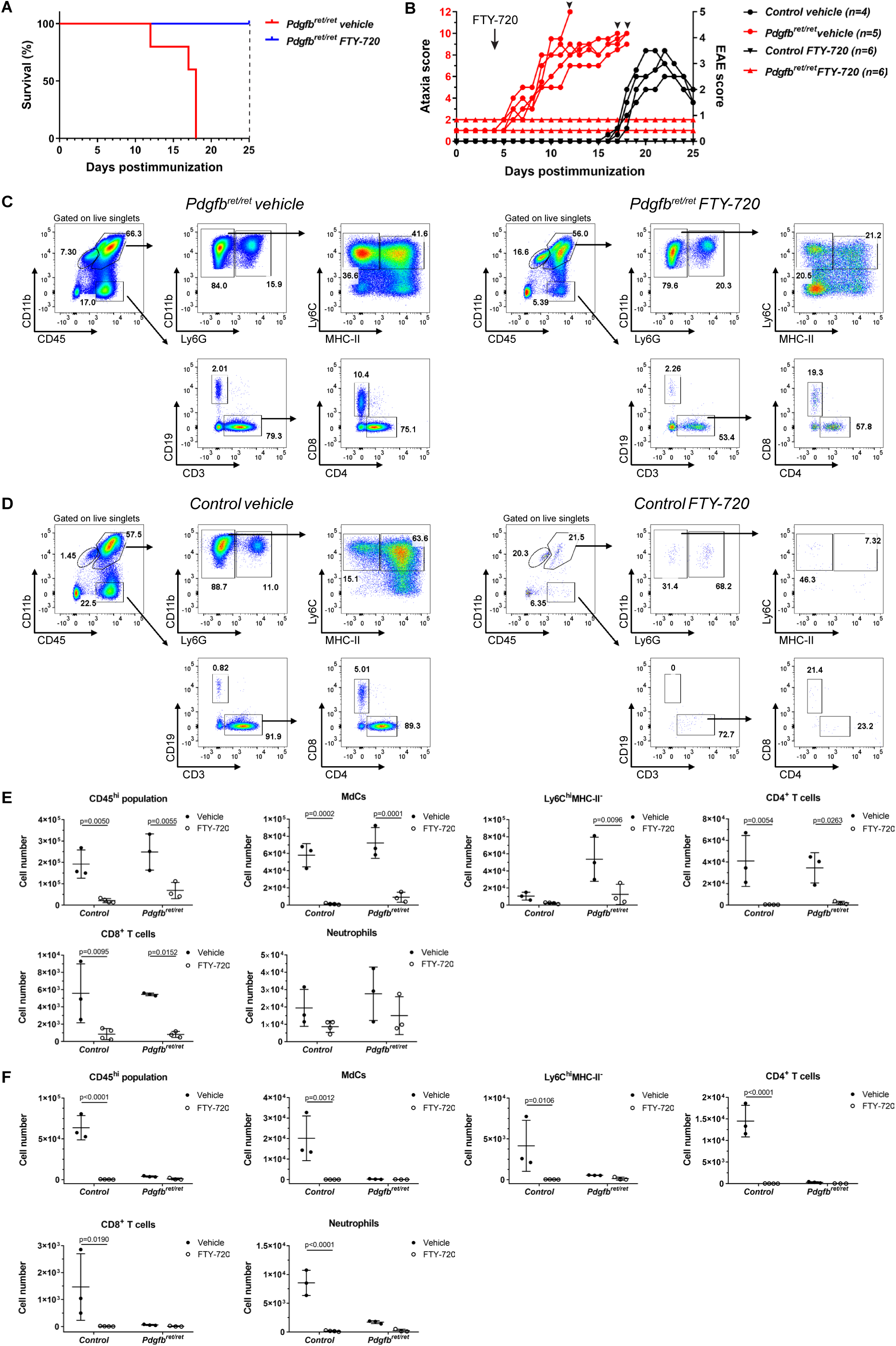
FTY-720 treatment rescues the lethality of pericyte-deficient mice after induction of EAE. (**a**) Kaplan-Meier survival curves of vehicle and FTY-720 treated *Pdgfb^ret/ret^* mice after active induction of EAE. The experiment was terminated on day 25 (marked by a black dashed line). n=5 mice treated with vehicle, n=6 mice treated with FTY-720. (**b**) Scoring of clinical symptoms during the course of FTY-720 treatment after induction of active EAE of control and *Pdgfb^ret/ret^* mice. The left y axis shows the cerebellar ataxia scores of *Pdgfb^ret/ret^* mice (in red) and the right y axis the classical EAE scores of control mice (in black). FTY-720 administration (0.5 mg/kg) was started on day 4 postimmunization (red dashed line) and the experiment was terminated on day 25 postimmunization. The ataxia score or EAE score of each mouse is plotted individually. Arrowheads indicate when *Pdgfb^ret/ret^* mice reached termination criteria and were sacrificed. N=4-5 mice per group. Representative flow cytometry pseudocolor plots showing the manual gating of microglia and other immune cell populations in the brains of vehicle or FTY-720 treated *Pdgfb^ret/ret^* mice (**c**) and in the spinal cords of vehicle or FTY- 720 treated control mice (**d**) after active induction of EAE. *Pdgfb^ret/ret^* mice were sacrificed for immune cell analysis when reached the ataxia score (8.5-10) and control mice were sacrificed for analysis when they reached EAE score 3-3.5. Quantification of the absolute cell numbers of the different immune cell populations (gating shown in c and d) in the brain (**e**) and spinal cords (**f**) of vehicle and FTY-720 treated control and *Pdgfb^ret/ret^* mice using flow cytometry. Controls: n=3 vehicle treated and n=4 FTY-720 treated, *Pdgfb^ret/ret^* mice: n=3 vehicle treated and n=3 FTY-720 treated. Data are presented as the mean ± SD. Two-way ANOVA test was used to determine statistical significance between groups.

### Spontaneous neuroinflammation in *Pdgfb^ret/ret^* mice expressing myelin specific T cell receptor

Little is known about what triggers spontaneous activation and entry of self-reactive T cells into the CNS. We asked whether the leukocyte permissive NVU in pericyte-deficient animals leads to spontaneous neuroinflammation when there is an overabundance of self-reactive T-cells towards a myelin antigen. To answer this question, we crossed *Pdgfb^ret/ret^* mice with 2D2 mice, which express a MOG_35-55_ peptide specific T cell receptor (TCR) ^21^. Previous studies have reported that approximately 5 % of 2D2 mice develop spontaneous EAE with classical symptoms ^21^. Offspring of *Pdgfb^ret/ret^* and *Pdgfb^wt/ret^;* 2D2*^tg^* crosses were monitored after weaning for signs of cerebellar ataxia and classical EAE. Similarly to previous observations (Fig. 4b, 5b), all animals carrying two alleles of mutated *Pdgfb* (*Pdgfb^ret/ret^*) presented with an ataxia score 2, consisting of hindlimb clasping (score 1) and kyphosis (score 1) already at weaning, which remained stable (Fig. 6a). However, *Pdgfb^ret/ret^; 2D2^tg^* mice showed increasing cerebellar ataxia scores compared to *Pdgfb^ret/ret^; 2D2^neg^* mice (Fig. 6a). Other control mice (*Pdgfb^wt/ret^; 2D2^neg^, Pdgfb^wt/ret^; 2D2^tg^*) occasionally received score 1, which was based on single balance loss on the ledge test. Of note, the ataxia score of individual *Pdgfb^ret/ret^; 2D2^tg^* mice fluctuated over the monitored time-period. None of the mice developed signs of classical EAE. Immunofluorescent staining of the brains of 3 months old *Pdgfb^ret/ret^; 2D2^tg^* animals showed an increased number of CD45^hi^ positive cells in the brains compared to *Pdgfb^ret/ret^; 2D2^neg^* mice (Fig. 6b). Flow cytometry of the immune cells confirmed that the brains of *Pdgfb^ret/ret^; 2D2^tg^* mice contain a significantly higher number of CD45^hi^ cells compared to *Pdgfb^ret/ret^; 2D2^neg^* (Fig. 6c, d). *Pdgfb^ret/ret^; 2D2^tg^* animals and all control animals were sacrificed for flow cytometry when *Pdgfb^ret/ret^; 2D2^tg^* animals had reached the ataxia score of 6-9. Interestingly, immune cell infiltrates of *Pdgfb^ret/ret^* and *Pdgfb^ret/ret^; 2D2^tg^* consisted of MdCs, Ly6C^hi^MHC-II^−^, CD4^+^ and CD8^+^ T cells (Fig. 2e and 6d). In the brains of *Pdgfb^ret/ret^; 2D2^tg^* mice, in addition to the aforementioned populations, neutrophils and B cells were detected (Fig. 6d). Spinal cords of *Pdgfb^ret/ret^; 2D2^tg^* mice did not show immune cell infiltrates (Supplementary Fig. 5) consistent with the *Pdgfb^ret/ret^* (Supplementary Fig. 1) or *Pdgfb^ret/ret^* mice during EAE (Fig. 4e, h). Thus, these experiments demonstrate that the leukocyte-permissive NVU caused by reduced pericyte coverage promotes the development of a neuroinflammatory disorder associated with increased myelin-reactive T cells in the circulation.

**Figure 6.**
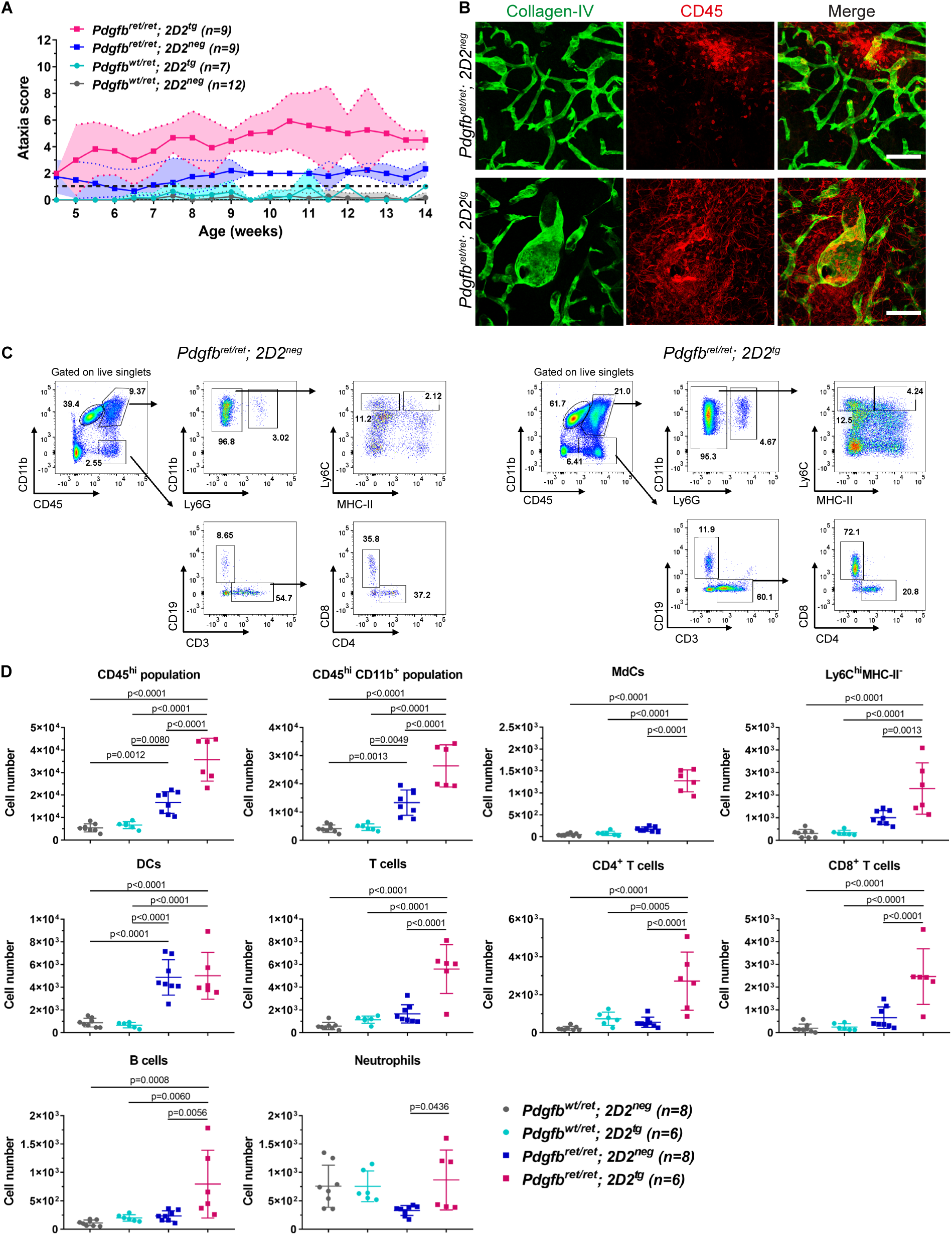
Pericyte-deficient mice expressing myelin specific T cell receptor develop neurological symptoms accompanied by immune cell infiltration. (**a**) Scoring of cerebellar ataxia in *Pdgfb^ret/ret^* x *Pdgfb^wt/ret^; 2D2^tg^* crossing offspring. Dashed black line indicates the mean baseline score of control mice (ataxia score=1, occasional slip during the ledge test). n=7-11 mice per genotype. (**b**) Immunofluorescent detection of CD45^hi^ leukocyte infiltrates and microglia (in red) in the striatum in *Pdgfb^ret/ret^; 2D2^neg^* and *Pdgfb^ret/ret^; 2D2^tg^* mice. Blood vessels are in green (collagen-IV). (**c**) Representative flow cytometry pseudocolor plots showing the manual gating of microglia and other immune cell populations in the brain. (**d**) Quantification of absolute cell numbers of immune cells (gated as shown in c) in the brains of control mice and *Pdgfb^ret/ret^; 2D2^neg^*, and *Pdgfb^ret/ret^; 2D2^tg^* animals. *Pdgfb^ret/ret^; 2D2^tg^* mice were terminated at the peak of cerebellar ataxia symptoms (ataxia score 6-9). Age of animals 2-3 months (**b, c**). Data are presented as the mean ± SD. Ordinary one-way ANOVA test was used to determine the statistical significance between groups. Scale bars are - 100 µm.

## Discussion

Pericytes have been shown to regulate BBB integrity at the level of endothelial transcytosis ^1, 2^. Pericytes also induce polarization of astrocyte end-feet ^1^; however, the extent of pericyte control over other characteristics of the brain vasculature is less explored. In this study, we investigated the role of pericytes in regulating leukocyte trafficking into adult CNS. In addition, we show that in the absence of pericytes, the NVU becomes permissive to leukocyte entry, leading to aggravated neuroinflammation in a setting of autoimmunity.

A previous study on pericyte-deficient *Pdgfb^−/−^* embryos showed that several LAMs, (e.g. *Icam1*, *Alcam*, *Lgals3*) were significantly upregulated on brain vasculature ^2^. In addition, a modest increase in Ly-6G/Ly-6C positive leukocytes was observed in the brains of juvenile *Pdgfb^F7/F7^* mice that display a 50% reduction in pericyte coverage compared to controls ^2^. Our observation that several LAMs, including VCAM-1 and ICAM-1, are upregulated in the adult vasculature of *Pdgfb^ret/ret^* mice (Fig. 1), which is accompanied by increased leukocyte entry into the brain parenchyma (Fig. 2), corroborates and extends these findings. Of note, endothelial cell-cell junctions in pericyte-deficient are closed to plasma proteins^1, 2^ indicating that increased leukocyte entry into the brain in pericyte-deficient mice is not due to relaxed cell-cell junctions. However, the molecular composition endothelial cell-cell contacts might be altered, which might facilitate leukocyte transmigration.

Similar persistent inflammation in the absence of pericytes as in the brain vasculature in *Pdgfb^ret/ret^* mice has been described in the retina ^22–25^. Whether an acute drop-out of pericytes in the adult organism leads to altered BBB permeability and alters the permissiveness of vasculature to leukocyte trafficking needs further studies. Pericyte ablation in adult mice using the *Pdgfrb*-Cre-Er^T2^; DTA mice was reported not to cause immediate BBB permeability changes in the retina and brain ^24^; however, pericyte loss in the brain was not assessed. Our unpublished data show that using the *Pdgfrb*-Cre-Er^T2^; DTA approach is not effective to deplete adult brain pericytes. In fact, only partial (up to 50 %) pericyte depletion could be achieved in the hippocampus. In adult vasculature, which has established structural (e.g. basement membrane) and cellular structures (e.g. astrocyte-end feet), pathological changes in the vasculature upon pericyte loss develop gradually over a few weeks. Thus, extensive loss of pericyte coverage during the formation of CNS vasculature results in sustained alterations at the NVU as well as at the level of vessel permeability and permissiveness to leukocyte trafficking, which cannot be corrected by other cellular constituents of the NVU. However, CNS vasculature compensates to a certain extent for pericyte-loss and reduced pericyte coverage. Upon a pericyte loss in the brain, the area covered by a single pericyte can be compensated by spatial rearrangement of nearby pericytes ^26^. Increased BBB permeability to intravenous tracers in juvenile *Pdgfb^F7/F7^* and *Pdgfb^F7/−^* mice that have a 50-60% reduction in pericyte coverage is absent in adult animals ^2^. The loss of pericytes may be compensated by other cells of the NVU such as astrocytes, which have been shown to promote BBB integrity and CNS immune quiescence ^27, 28^. Previous studies have shown an inverse correlation between pericyte coverage and BBB permeability in the brain ^1, 2, 14^. Based on previous data on pericyte-deficient mice, up to 50 % reduction in pericyte vessel coverage does not lead to overt BBB permeability changes (Fig. 7a). Notably, an increased infiltration of leukocytes into brain parenchyma has been reported in mouse pups that express one allele of constitutively active PDGFRB, which alter pericyte differentiation ^29^, indicating that both altered pericyte numbers/vessel coverage and activation state ^29, 30^ disturb pericyte-endothelial signaling. In the normal CNS, this disturbed signaling limits immune surveillance.

**Figure 7.**
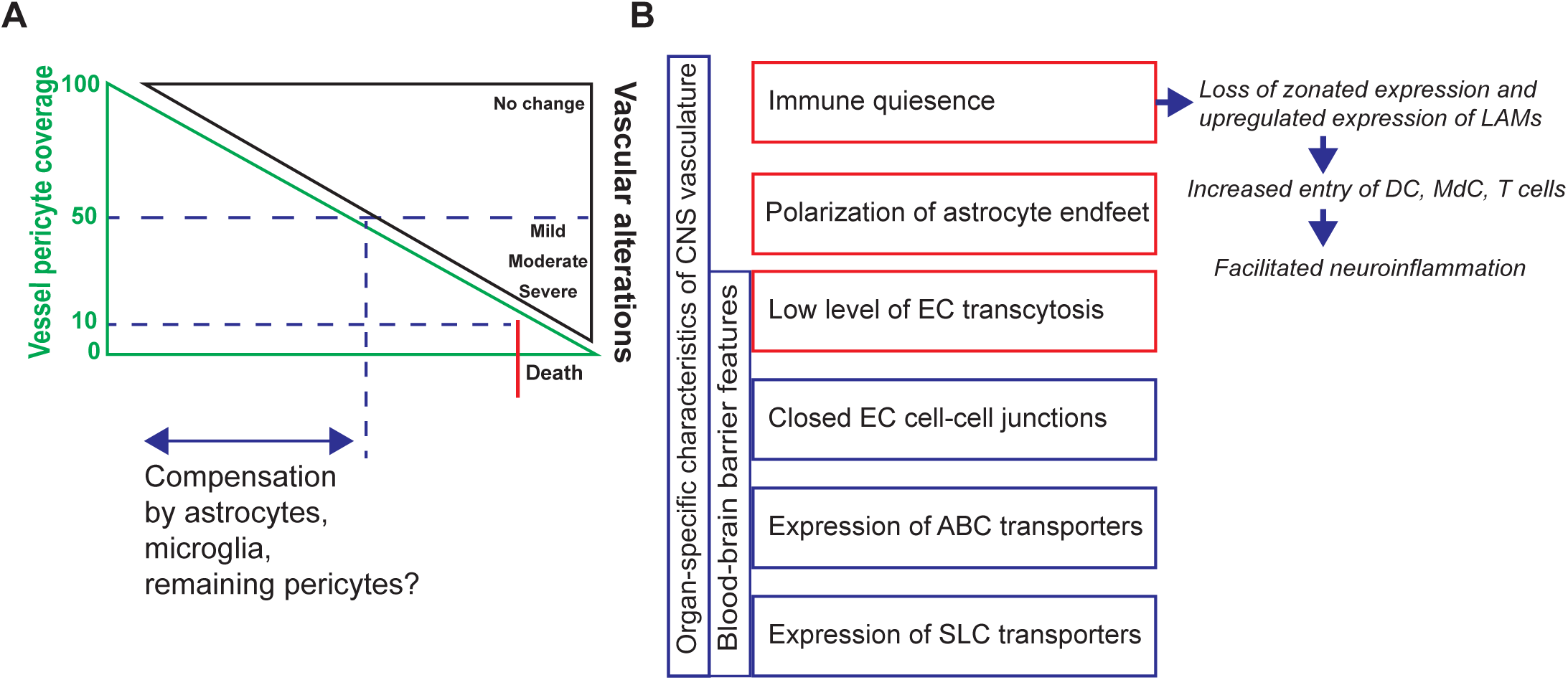
The consequences of pericyte-loss on CNS vasculature during development. (**a**) Reduction of vessel pericyte coverage correlates with the severity of BBB alterations. Up to 50-60 % of pericyte loss during the development does not result in overt BBB changes ^2^. Changes occurring at the NVU due to the reduced pericyte coverage could be compensated by remaining pericytes and/or by other cellular constituents of the NVU (e.g. astrocytes). Approximately 85% - 40% loss of pericytes is compatible with development; however, it results is persistent changes at the NVU, which cannot be compensated by other cellular components of the NVU. Pericyte coverage less than 10-15% during development results in premature death. (**b**) Pericytes regulate organ specific characteristics of the CNS vasculature. Previous studies have shown that pericytes regulate the BBB permeability on the level of endothelial transcytosis ^1, 2^ and polarization of astrocyte end-feet ^1^. Of note, pericytes do not regulate expression of ABC transporters ^1^. Accordingly, increased BBB permeability to plasma proteins is not paralleled by increased transport of small-molecular drugs ^67^. Our study corroborates pervious data showing an increased expression of LAMs upon the absence of pericytes ^2^. Poor pericyte coverage in adult mice results in the loss of zonated expression and upregulation of VCAM-1 and ICAM-1 on blood vessels which licences entry of dendritic cells (DC), monocyte-derived cells (MdC) and T cells into the CNS, which upon the induction of neuroinflammation aggravates the disease and determines the anatomical location of immune cell infiltrates.

Increased infiltration of immune cells into the brain parenchyma in adult *Pdgfb^ret/ret^* is not accompanied by demyelination. This finding indicates that the permissive state of brain vasculature to immune cell entry is not sufficient to trigger demyelinating CNS pathology. Normal oligodendrocyte differentiation and myelination of adult *Pdgfb^ret/ret^* mice has also been investigated ^31^. The findings of this study contradict a previous report of white matter changes and loss of myelin in another mouse model of pericyte-deficiency – *Pdgfrb^F7/F7^* mice ^32^. However, several other studies have not found evidence that adult *Pdgfrb* mutants demonstrate pericyte loss during aging or present altered BBB permeability ^2, 14^. Whereas pericytes and pericyte-like cells have been shown to proliferate and promote oligodendrocyte differentiation during remyelination after acute CNS injury ^31^, mechanisms other than pericyte loss in old *Pdgfrb* mutants could contribute to the reported white matter changes during aging (i.e. dysfunction in neural cell types other than pericytes expressing PDGFRB).

Interestingly, we observed an increased number of CD11c^+^ DCs in the brain parenchyma of adult *Pdgfb^ret/ret^* mice. The presentation of CNS antigens by DCs leads to activation of autoreactive CD4^+^ helper T cells in CNS parenchyma, which is a crucial step for licensing T-cells to initiate neuroinflammation ^33–35^. The CD11c^+^ DCs have been shown to interact and transmigrate across inflamed CNS endothelium in an integrin α4β1dependent manner during EAE ^36^. However, the molecular mechanisms of DCs trafficking to the CNS under homeostasis is less well understood ^37^. Thus, the increased number of DCs in the brain parenchyma of *Pdgfb^ret/ret^* mice indicates that intrinsic changes of the NVU due to reduced pericyte coverage promote immune surveillance, which could facilitate the initiation of neuroinflammation observed during EAE (Fig. 4, Supplementary fig. 4) and pericyte-deficient animals expressing MOG_35-55_ specific TCR (Fig. 6 and Supplementary fig. 6, Fig. 7b).

Brain-restricted neuroinflammation in pericyte-deficient mice is dominated by myeloid cells (CD11b^+^Ly6C^hi^) – approx. 87 % and 17 % of the parent population (CD45^hi^CD11b^+^Ly6G^−^) in *Pdgfb^ret/ret^* mice during EAE and in *Pdgfb^ret/ret^*;*2D2^tg^* mice, respectively (Fig. 4f, g and Fig. 6c, d). Increasing evidence points to the key role of myeloid cell subsets in mediating tissue damage in EAE and MS (reviewed in ^38, 39^). Infiltration of CCR2^+^Ly6C^hi^ inflammatory monocytes from the blood into the CNS parenchyma coincides with the onset of the clinical signs and worsens the severity of EAE ^40–42 43^.

The localization of demyelinating lesions in MS patients shows a variable pattern. Typical affected areas are cerebral white matter (e.g. periventricular, corpus callosum), brain stem and cerebellum ^44^ as well as optic nerve and spinal cord. In the case of neuromyelitis optica spectrum disorders, another type of autoimmune demyelinating disease, the optic nerve and spinal cord are preferentially damaged. ^45^. In EAE, a “classical” spinal cord phenotype (ascending flaccid paralysis) can be distinguished from “atypical” EAE with inflammation localized to the cerebrum, cerebellum and brain stem. The underlying mechanisms leading to regional differences in leukocyte extravasation are not well understood. It has been suggested that brain and spinal cord are distinct microenvironments with a distinct inflammatory cell repertoire, including different T cell types and different cytokines ^46^. We found that upon induction of autoreactive neuroinflammation, *Pdgfb^ret/ret^*, mice succumb to atypical EAE phenotype (Fig. 4b). In addition, *Pdgfb^ret/ret^*; *2D2^tg^* animals presented fluctuating symptoms of cerebellar ataxia (Fig. 6a), milder than after the induction of EAE. Histological and flow cytometry analyses of infiltrated immune cells, which showed neuroinflammation localized to the brain in pericyte-deficient mice, were in concordance with the clinical deficits (Fig. 4 and Fig. 6). We showed that although the vessel pericyte-coverage is reduced in the spinal cord of *Pdgfb^ret/ret^* mice compared to controls (Fig. 3b), pericyte vessel-coverage in the spinal cord in *Pdgfb^ret/ret^* mice is more complete compared to brain ^1, 14^. This relatively spared vasculature of spinal cord vessels with a higher pericyte-coverage and lack of upregulation of ICAM-1 and VCAM-1 could explain the preferential location of neuroinflammation in the brain in *Pdgfb^ret/ret^* and *Pdgfb^ret/ret^*; *2D2^tg^* mice (Fig. 7).

BBB breakdown, one of the pathological hallmarks of MS ^47, 48^, is an early event in the formation of the inflammatory lesions and has been suggested to precede parenchymal inflammation ^49^. Interestingly, one of the early changes after induction of EAE, at the capillary level, increased transcytosis of brain endothelial cells causing increased vessel permeability ^50^. Pericyte damage (e.g. lipofuscin accumulation, membrane protrusions) in chronic-progressive has been described in MS lesions^51^. It has been suggested that the BBB becomes disrupted by early inflammatory microlesions via IL-1β ^52^ produced my MdCs and neutrophils after EAE induction ^53^. However, the nature of changes at the NVU that precede immune cell entry and cause failure of vascular the immune regulatory function in MS is not known. Our study shows that intrinsic changes in brain vasculature facilitate the neuroinflammatory cascade and can influence the localization of the neuroinflammatory lesions. In the future, it would be interesting to investigate whether intrinsic changes in brain vasculature such as alterations in pericyte-endothelial cross talk leading to a pro-inflammatory profile of endothelial cells regulate the localization of MS lesions.

In conclusion, our study demonstrates that pericytes contribute to CNS immune quiescence by limiting leukocyte infiltration into the CNS during homeostasis and autoreactive neuroinflammation. In addition, immune cells preferentially home into CNS regions in which vessels show a proinflammatory profile due to the dramatically reduced pericyte coverage. The presence of abundant myelin peptide-specific peripheral T cells is sufficient to promote the development of spontaneous autoimmune brain inflammation. Future studies should be aimed at unraveling the molecular mechanism leading to vascular permissiveness to leukocyte entry in the setting of altered pericyte coverage or/and pericyte-endothelial cross talk. Since vascular dysfunction modulates leukocyte entry and neuroinflammation, vasoprotective therapies combined with pre-existing treatments could lead to improved clinical outcome in MS.

## Methods

### Mice

Mice were kept in individually ventilated cages under specific-pathogen-free conditions. The following genetically modified mouse lines were used for experiments: PDGFB-retention motif knock out (*Pdgfb^ret/ret^*) ^22^ and myelin oligodendrocyte glycoprotein (MOG_35-55_) specific T cell receptor transgenic mouse line (*2D2*) ^21^. Mice were kept on a C57BL/6J genetic background. Animal experiment protocols were approved by the Veterinary office of the Canton of Zurich (permits ZH196/204, ZH070/2015, ZH151/2017, ZH072/2018).

### Histochemistry and immunohistochemistry (IHC)

Mice were transcardially perfused under anesthesia with Hank’s balanced salt solution (HBSS) and 4% paraformaldehyde (PFA) subsequently. The organs of interest were dissected, postfixed in 4% PFA overnight and embedded into paraffin. The stainings were performed on 2 µm thick sections, except for the luxol fast blue – periodic acid Schiff (LFB-PAS) staining, which were 5 µm thick. LFB-PAS staining was performed according to standard procedure. Deparaffinized and rehydrated sections were incubated in luxol-blue solution for 30 min at 60-70 °C, differentiated in lithium-carbonate solution and counterstained with 1% periodic-acid solution for 10 min, Schiff-reagent for 25 min and hematoxylin for 1 min. CD3 and B220 immunostaining were performed according to standard procedure using an Automated IHC Stainer (Bond-III, Leica Biosystems). Deparaffinized and rehydrated sections were incubated with CD3 (Thermo Fisher Scientific, cat. # RM-9107-s, 1:50) or B220 (BD Biosciences, cat. # 553084, 1:8000) for 30 min respectively, subsequently incubated with Bond Polymer Refine Red Detection solution (with alkaline phosphatase (AP), Leica Biosystems) for 30 min or with Bond Polymer Refine Detection solution (with 3,3’-Diaminobenzidine (DAB), Leica Biosystems) and hematoxylin for 10 min. For CD3 staining, the antigen retrieval with EDTA (pH 8) was performed for 20 min. Stained paraffin sections were scanned with NanoZoomer Digital Pathology (Hamamatsu Photonics).

### Immunofluorescent stainings

Mice were transcardially perfused under anesthesia with HBSS and then 4% PFA. Brains and spinal cords were dissected and postfixed for 4-6 hours in 4% PFA. 60 µm thick vibratome sections were incubated in blocking and permeabilization buffer (1% BSA, 2% Triton-X in PBS) overnight at 4°C. Subsequently, sections were incubated with primary antibody mix for 2 days overnight at 4°C, afterwards with secondary antibody mix for 1 day overnight at 4°C and finally with DAPI (1:10000, Sigma-Aldrich) for 8 min at room temperature. Stained sections were mounted with Mount Prolong Gold Antifade Mountant (Invitrogen). The following primary antibodies were used: rabbit anti-mouse collagen-IV (Bio-Rad Company, cat.# 2150-1470, 1:300), rat anti-mouse CD45 (BD Pharmingen, cat.# 553076, 1:100), rat anti-mouse VCAM-1 (Merck, cat.# CBL-1300, 1:100), rat anti-mouse ICAM-1 (Abcam, cat.# ab119871, 1:100), goat anti-mouse CD13 (R&D Systems, cat.# AF2335, 1:100) and chicken anti-mouse MBP (Millipore, cat.# AB9348, 1:100). Fluorescently labelled secondary antibodies suitable for multiple labelling were purchased from Jackson Immuno Research. Images were taken by Leica SP5 confocal laser scanning microscope (Leica microsystems, 20X objective, NA=0.7). Images were analyzed by the image processing software Fiji and Imaris. Images were postprocessed by Adobe Photoshop and Adobe Illustrator.

### Quantification of MBP immunofluorescence

Images of immunofluorescently labeled sections were acquired by Leica SP5 confocal laser scanning microscope (Leica microsystems, 20X objective, NA=0.7) equipped with hybrid detectors (HyD) in photoncounting mode. Mean fluorescent intensity was calculated in three regions of interests (ROIs) of the corpus callosum in coronal brain sections using the mean grey value measurement tool in Fiji. The average value of fluorescent intensity per animal was calculated and plotted on a graph.

### Quantification of vessel pericyte, VCAM-1 and ICAM-1 coverage

Images of immunofluorescently labeled sections were acquired by Leica SP5 confocal laser scanning microscope (Leica microsystems, 20X objective, NA=0.7) equipped with HyD. Pericyte coverage was calculated using the area measurement tool in Fiji. The area of CD13 and collagen-IV signal was measured on binary images in 6 ROIs, 150 x 150 µm each, in coronal spinal cord sections. Coverage was calculated as the percentage of CD13 positive area over the collagen-IV positive area. The collagen-IV area was taken arbitrarily as 100% and the CD13 positive area was expressed as a percentage normalized to the collagen-IV area. VCAM-1 and ICAM-1 vessel coverage was calculated using the area measurement tool in Fiji. The area of VCAM-1 or ICAM-1 and the GLUT-1 signal (on VCAM-1 stained sections) or collagen-IV signal (on ICAM-1 stained sections) was measured on binary images in 6 ROIs, 200 x 200 µm each, in coronal brain sections. Coverage was calculated as the percentage of VCAM-1 or ICAM-1 positive area over the GLUT-1 or collagen-IV positive area, respectively. The GLUT-1 or collagen-IV area was taken arbitrarily as 100% and the VCAM-1 or ICAM-1 positive area was expressed as a percentage normalized to GLUT-1 or collagen-IV area, respectively.

### Quantification of CD45^hi^ cell infiltrates

Images of immunofluorescently labeled sections were acquired by Leica SP5 confocal laser scanning microscope (Leica microsystems, 20X objective, NA=0.7) equipped with photomultiplier tube (PMT) detectors. Two ROIs were analysed in 24 sections in the cortex, in 19 sections in the striatum and in 15 sections in the corpus callosum on horizontal brain sections. A surface mask was added to the collagen-IV signal and the CD45 signal was marked using the “spots” function in each ROI using Imaris x64 software. The total number of spots (CD45^hi^ cells) and number of spots not overlapping with the collagen-IV surface mask (extravasated CD45^hi^ cells) was calculated using the built-in statistics tool of Imaris x64 software. The percentage of spots not overlapping with the collagen-IV surface mask to the total number of spots was calculated per each ROI. Finally, the average percentage of spots far from the collagen-IV surface mask was calculated per brain region and per animal and plotted on a graph.

### Transmission electron microscopy

Mice were transcardially perfused under anesthesia with HBSS followed by 2% PFA, 2.5% glutaraldehyde in 0.1 M cacodylate buffer (pH 7.4). Brains were dissected, 1 mm thick coronal sections were cut with a brain matrix (World Precision Instruments, cat.# RBMA-200C) and kept in 2% PFA, 2.5% glutaraldehyde in 0.1 M cacodylate buffer (pH 7.4) until sample preparation. 1 mm^3^ thick blocks were cut manually from the region of corpus callosum, washed in 0.1 M cacodylate buffer (pH 7.4) and incubated in 1% osmium-tetroxide in 0.1 M cacodylate buffer (pH 7.4) for 1 hour. After two washing steps in distilled water, the tissue blocks were contrasted with uranyl-acetate in distilled water overnight, followed by dehydration with a series of alcohol and embedded in Epon resin. Ultrathin sections (70 nm) were contrasted with lead-citrate and mounted on grids for imaging. Imaging was performed on FEI CM100 transmission electron microscope (Philips) using an Orius 1000 digital CCD camera (Gatan, Munich, Germany).

### Scoring of demyelination on LFB-PAS stained sections

Three LFB-PAS stained sections per animal were assessed for myelination. The scoring system to assess demyelination was adopted from Kim et al. ^54^. Demyelination scores for the medial corpus callosum were subjectively evaluated as follows: 0 - fully myelinated; 1 - mild demyelination (≤1/3 affected) in the center; 2 - moderate demyelination (≤2/3 affected) in the center; 3 - no myelin in the center; 4 - demyelination extending to the arch of the medial corpus callosum. Demyelination scores for lateral projections of the corpus callosum were scored as follows: a range from 0 to 3 was used, where 0 equals to normal myelination and 3 indicates no myelin.

### Leukocyte and microglia isolation from the CNS

Mice were transcardially perfused under anesthesia with HBSS. Brains and spinal cords were dissected and digested with collagenase-D (analysis of EAE and 2D2 experiments, Sigma-Aldrich) or collagenase Type IV (Sigma Aldrich) in Roswell Park Memorial Institute medium (RPMI) 1640. The digested tissue was washed with PBS and centrifuged. The CNS pellet was subsequently centrifuged in 30% Percoll (GE Healthcare) gradient at 10800 rpm for 30 min (analysis of EAE and 2D2 experiments) or 1560 rpm for 20 min (analysis of naïve mice) to remove myelin. After myelin removal the single cell suspension was washed with PBS, centrifuged and resuspended in FACS buffer (2% fetal bovine serum (FBS), 0.01% NaN_3_ in PBS) for flow cytometry analysis.

### Leukocyte isolation from the lymph nodes, spleen, thymus and blood

200 µl blood was drawn before mice were transcardially perfused under anesthesia with HBSS. Lymph nodes, spleen, thymus were dissected. Four lymph nodes (axillary and inguinal) were measured per mouse. Organs were filtered through a 70 µm mesh. Spleen and blood samples were lysed with red blood cell lysis buffer (150 mM NH_4_Cl, 10 mM KHCO_3_, 0.1 mM Na_2_EDTA, pH 7.4). The single cell suspension was washed with PBS and resuspended in FACS buffer for flow cytometry analysis. The total cell number was counted with an automated cell counter (BioRad, TC20).

### Flow cytometry analysis

For cell surface staining, cells were incubated with mix of fluorophore-labelled primary antibodies for 30 min on ice in the dark. Cells were resuspended in FACS buffer. Primary antibodies were purchased from Biolegend and BD Biosciences and are listed in Supplementary Table 1) Flow cytometry analysis was performed on LSR II Fortessa (Becton Dickinson) and data were analyzed with FlowJo (v10.2 and v10.5.3) software.

### Automated population identification in high-dimensional data analysis

Raw data was pre-processed using FlowJo followed by transformation in Matlab using cyt2, normalization in R obtaining values between 0 and 1, automated and unsupervised two-dimensional cell mapping, dimensionality reduction and visualization by t-distributed stochastic neighbour embedding (t-SNE) ^55, 56^. The FlowSOM algorithm was used for automated clustering ^57^ using the t-SNE map with overlaid mean marker expression values and a heatmap of median expression values ^56, 58^. Frequencies for each cluster was identified using R, exported into an excel file and used for further analysis.

### Induction of EAE

For active immunization ^59^ mice were immunized subcutaneously into the flanks with 200 µg MOG_35-55_ peptide (Anawa) emulsified in complete Freund’s adjuvant (CFA; InvivoGen) and 200 ng pertussis toxin (List Biological Laboratories Inc.) intraperitoneally on day 0 and day 2 post-immunization. For passive immunization (adoptive transfer) ^60^ experiments C57BL/6J mice (Janvier Labs) were immunized as described above omitting the injection of pertussis toxin on day 2 post-immunization. On day 7, splenocytes and lymphocytes from the draining lymph nodes were isolated and cultured for 2 days under polarizing conditions (20 µg/ml MOG_35-55_, 10 ng/ml IL-23). Recipient mice (control and *Pdgfb^ret/ret^*) were sublethally irradiated with 550 rad. One day after irradiation, recipient mice received intraperitoneally 11 x 10^6^ MOG-activated cells. Control or *Pdgfb^ret/ret^* mice were terminated for histology or flow cytometry analysis when they reached termination criteria - classical EAE score of 3-3.5 or an ataxia score of 8.5-10, respectively.

### FTY-720 treatment and termination criteria for immune cell analysis

Mice were treated daily by oral gavage with 0.5 mg/kg FTY-720 (Selleckhem) in 2% (2-hydoxypropyl)-β-cyclodextrin (Sigma-Aldrich) starting on day four post-immunization. Controls groups received 2% (2-hydoxypropyl)-β-cyclodextrin in water (vehicle). Vehicle treated mice were terminated for histology or flow cytometry analysis when control or *Pdgfb^ret/ret^* mice reached a classical EAE score of 3-3.5 or an ataxia score of 8.5-10, respectively. FTY-720 treated mice were terminated on the same day when vehicle treated mice reached termination criteria.

### Scoring to assess the severity of EAE

Mice were monitored daily after induction of EAE. The scoring scale of *classical EAE* symptoms: 0 - no detectable clinical signs; 1 - complete tail paralysis; 2 - unilateral partial hind limb paralysis; 2.5 - bilateral partial hind limb paralysis; 3 - complete bilateral hind limb paralysis; 3.5 - complete bilateral hind limb and partial forelimb paralysis; 4 - moribund, complete fore- and hind limb paralysis; 5 - dead.

The scoring of *atypical EAE* (cerebellar ataxia) symptoms was adopted from Guyenet et al. ^18^. Following four individual parameters were scored - ledge test, hind limb clasping, gait and kyphosis. Each individual measure was scored on a scale from 0 to 3 and recorded at the end of the measurement as a combined score from 0 to 12. The scale for *ledge test* was assessed as follows: 0 - the mouse is able to walk along the ledge without losing its balance and lands on its paws when lowering itself back to the cage; 1 - the mouse is not able to walk along the ledge without losing its footing, but otherwise walks coordinated; 2 - the mouse is not able to effectively use its hind limbs while walking along the ledge and lands rather on its head then its paws; 3 -: the mouse is completely unable to walk along the ledge or falls off. The scale for the *hind limb clasping* was assessed as follows: 0-upon lifting the mouse by its tail both of its hind limbs are persistently pointing away from the abdomen; 1 - if one hind limb is pulled back towards the abdomen for more than 50% of the time suspended; 2 - both hind limbs are partially pulled back towards the abdomen for more than 50% of the time suspended; 3 - hind limbs are entirely retracted and touching the abdomen for more than 50% of the time. The scale for *gait* was assessment is as follows: 0 - the mouse walks coordinated, all four limbs support the body weight evenly and its abdomen is not touching the ground; 1 - the mouse limps while walking or tremor can be observed; 2 - the mouse has severe tremor or severe limp or lowered pelvis and the feet are pointing away from the body (“duck feet”); 3 - the mouse has severe difficulties to walk and the abdomen is completely touching the ground or it refuses to walk at all. The scale for *kyphosis* was assessed as follows: 0 - the mouse can freely straighten its spine while walking, no visible kyphosis; 1 - the mouse has a mild kyphosis but is able to straighten its spine while walking; 2 - the mouse has a constant kyphosis and is unable to straighten its spine completely; 3 - the mouse has a pronounced kyphosis while walking or sitting.

### Transcriptional profile analysis of microvasculature of pericyte deficient mice

*Pdgfb^ret/ret^* and control mice brain microvascular transcriptome data have been published previously (GSE15892) ^1^. Normalized intensity values and meta data were obtained from Gene Expression Omnibus, using the “GEOquery#x201D; package from Bioconductor ^61^. Probe annotation data were obtained using the “mouse4302.db” package, also from Bioconductor ^62^. Probe-level intensity values were summarized at the gene-level using the collapseRows functionality within the WGCNA R package ^63^. If a gene symbol was associated with more than one probe id, the probe id showing maximum variation was kept for downstream analyses. Differential expression summary statistics were computed using the empirical Bayes approach implemented with the “limma” package from Bioconductor ^64^. Heatmap visualizing the expression pattern of selected genes was created using the “pheatmap” R package ^65^. All bioinformatics analyses were performed in R version 3.5.1 ^66^.

### Statistical analysis

Statistical significance was determined with the unpaired t-test or ANOVA (GraphPad Prism 8.0). Differences with a p value <0.05 were considered statistically significant. Data is presented as the mean ± SD.

## Author contribution

The conception of the study (B.S., A.K.), the design of the work (O.T., B.S.; M.T.H., M.G., B.B., A.K.); the acquisition of data (O.T., B.S., H-C.T., J.S., A.K.), all authors contributed to the analysis or interpretation of data. O.T. and A.K. wrote the manuscript with substantial input from B.S., S.U., M.H.H., B.B., M.G.. All authors reviewed the manuscript.

## Acknowledgements

Samples for electron microscopic imaging were prepared by the Center for Microscopy and Image Analysis of the University of Zürich. Imaging was performed with equipment maintained by the Center for Microscopy and Image Analysis of the University of Zürich. Flow cytometry analysis was performed with equipment maintained by the Flow Cytometry Facility of the University of Zürich. This work was supported by grants from Swiss National Science Foundation (grant 31003A_159514), Swiss Heart Foundation, The Swiss Cancer League (grant KLS-3848-02-2016) to A.K., Swiss Society of Multiple Sclerosis to A.K. and B.S., and the Leducq Foundation (grant 14CVD02) to A.K. and M.H.H.

