## Supplemental data for "Pericytes regulate vascular immune homeostasis in the CNS"

**Supplementary Figure 1. Deregulated expression of LAMs in pericyte-deficient brain vasculature and the lack of CD45$^{hi}$ leukocyte infiltrates in the spinal cord of *Pdgfb$^{ret/ret}$* mice.** (**a**) The heatmap representing of relative gene expression of LAMs in microvascular fragments of *Pdgfb$^{ret/ret}$* mice compared to controls. The scale shows z-score of normalized intensities. (**b**) Immunofluorescent staining of the vasculature (collagen-IV, in green) and CD45$^{hi}$ leukocytes and microglia (in red) in the spinal cord of control and *Pdgfb$^{ret/ret}$* mice. (**c**) Representative flow cytometry pseudocolor plots showing manual gating of microglia and other immune cell populations in the spinal cord. (**d**) Quantification of the absolute cell numbers of different immune cell populations (gating shown in c) in the brains and spinal cords of naïve control and *Pdgfb$^{ret/ret}$* mice. Scale bars are 100 μm.

**Supplementary Figure 2. Flow cytometry and histological analysis of primary and secondary lymphoid organs and blood.** (**a**) Total cell numbers in the blood, thymus, spleen and lymph nodes analysed with flow cytometry. N=3 mice per genotype. Leukocyte subpopulations in the blood (**b**), spleens (**c**) and lymph nodes (**d**) measured with flow cytometry. N=3 mice per genotype. Haematoxylin-eosin (HE), CD3 (in red) and B220 (in red) stainings visualizing the structure of lymph nodes (**e**), thymuses (**f**) and spleens (**g**) of 2-3 months old adult control and *Pdgfb$^{ret/ret}$* mice. CD3 and B220 immunostainings were counterstained with haematoxylin to visualize cell nuclei. Data are presented as the mean ± SD. Scale bars: e, g - 300 μm, f - 100 μm.

**Supplementary Figure 3. Assessment of myelination in the corpus callosum in naïve control and *Pdgfb$^{ret/ret}$* mice.** (**a**) Images of LFB-PAS stained brain sections showing the medial corpus callosum. N= 6 control mice, n=3 *Pdgfb$^{ret/ret}$* mice. (**b**) Quantification of demyelination in the medial and lateral corpus callosum (cc). N= 3 mice per genotype. Two-way ANOVA test was used to determine statistical significance between groups. Image of the coronal brain section is from the Allen Mouse Brain Atlas. (**c**) Immunofluorescent images showing expression of myelin basic protein (MBP) in the brain of 9-11 months old naïve mice. The image of the coronal brain section is from the Allen Mouse Brain Atlas. (**d**) Quantification of MBP fluorescent intensity in the medial corpus callosum of young adult mice (2-3 months old). Unpaired t-test was used to determine the statistical significance (**e**) Electron microscopic images showing myelination in the corpus callosum of young adult mice (2-3 months old). Data are presented as the mean ± SD. Scale bars are: a - 300 µm, b – 2 mm, c - 2 µm.

**Supplementary Figure 4. Histological and cellular changes after induction of active EAE in the CNS.** (**a**) Kaplan-Meier survival curves of *Pdgfb*$^{wt/wt}$, *Pdgfb*$^{wt/ret}$ and *Pdgfb*$^{ret/ret}$ mice after active induction of EAE. (**b**) EAE score of *Pdgfb*$^{wt/wt}$ and *Pdgfb*$^{wt/ret}$ mice did not show any difference in the course of the disease. (**c**) Immunofluorescent staining showing CD45$^{hi}$ infiltrated leukocytes and microglia (in red) in the brains of control (EAE score 3) and *Pdgfb*$^{ret/ret}$ (ataxia score 9) mice after active EAE. Blood vessels are in green (collagen-IV) and myelin in cyan (MBP). The yellow dotted lines surround ventricles (marked with *). (**d**) Infiltrated T cells (CD3, in brown) in the cortex and corpus callosum of *Pdgfb*$^{ret/ret}$ (day 11 pi) mice. CD3 immunostainings were counterstained with haematoxylin visualizing cell nuclei. (**e**) Images of LFB-PAS stained brain sections showing the medial corpus callosum after induction of active EAE. (**f**) Demyelination scores in the medial and lateral corpus callosum of (cc) control (EAE score 3) and *Pdgfb*$^{ret/ret}$. Two-way ANOVA test was used to determine statistical significance between groups. N= 7 control mice, n=6 *Pdgfb*$^{ret/ret}$ mice. Quantification of absolute cell numbers of neutrophils and B cells (gated as shown in Fig. 3f and h) in the brains (**g**) and spinal cords (**h**) of control and *Pdgfb*$^{ret/ret}$ mice using flow cytometry. Pooled data from two individual experiments. N= 5 mice per genotype. Data are presented as the mean ± SD. Scale bars are: c - 300 µm, d - overview image: 1 mm, cortex, 1, 2, 1': 100 µm, 3, 2': 10 µm.

**Supplementary Figure 5. Analysis of the blood and brain tissue after FTY-720 treatment.** (**a**) Representative flow cytometry pseudocolor plots showing manual gating of $CD4^+$, $CD8^+$ T cell and B cell populations in the blood on day 12 post-immunization with MOG peptide. Quantification of the leukocyte (**b**) and T cell (**c**) numbers per 50 µl blood (gated as in a) with flow cytometry. (**d**) Immunohistochemical staining of CD3 positive T cells (in red) on sagittal brain cuts shows reduction of T-cell infiltrates the cerebellum and brain stem of $Pdgfb^{ret/ret}$ mice after FTY-720 treatment (day 16). Tissue sections were counterstained with hematoxylin (in blue). Scale bars: 100 µm. Data are presented as the mean ± SD. Two-way ANOVA test was used to determine the statistical significance between groups.

**Supplementary Figure 6. Analysis of immune cells in the spinal cord of pericyte-deficient mice expressing myelin specific T cell receptor.** (**a**) Representative flow cytometry pseudocolor plots showing manual gating of microglia and other immune cell populations in the spinal cord. (**d**) Quantification of absolute cell numbers of immune cells (gated as shown in a) in the spinal cords of control mice and $Pdgfb^{ret/ret}$*; 2D2*$^{neg}$, and $Pdgfb^{ret/ret}$*; 2D2*$^{tg}$ animals. $Pdgfb^{ret/ret}$*; 2D2*$^{tg}$ mice were terminated at the peak of the cerebellar ataxia symptoms (ataxia score 6-9). One-way ANOVA test was used to determine the statistical significance between groups.

**Supplementary Table 1. Antibodies used for flow cytometry**

| Antibody | Company | Catalogue number |
|---|---|---|
| anti-mouse CD45 – PE | Biolegend | 103125 |
| anti-mouse CD3 – FITC | Biolegend | 100203 |
| anti-mouse CD4 – BV711 | Biolegend | 100549 |
| anti-mouse CD8 – PE-CF594 | BD Pharmingen | 562315 |
| anti-mouse CD19 – PE | BD Pharmingen | 553786 |
| anti-mouse Ly6G – Alexa Fluor 647 | Biolegend | 127609 |
| anti-mouse MHC-II – Alexa Fluor 700 | Biolegend | 107621 |
| anti-mouse CD11c – PE-Cy7 | Biolegend | 117317 |
| anti-mouse CD11b – APC-Cy7 | BD Pharmingen | 561039 |
| anti-mouse Ly6C – BV605 | Biolegend | 128035 |
| anti-mouse CD19 – PerCP-Cyanine 5.5 | Biolegend | 115533 |
| anti-mouse Ly6G – Alexa Fluor 700 | Biolegend | 127621 |
| 7-AAD | BD Pharmingen | 559925 |

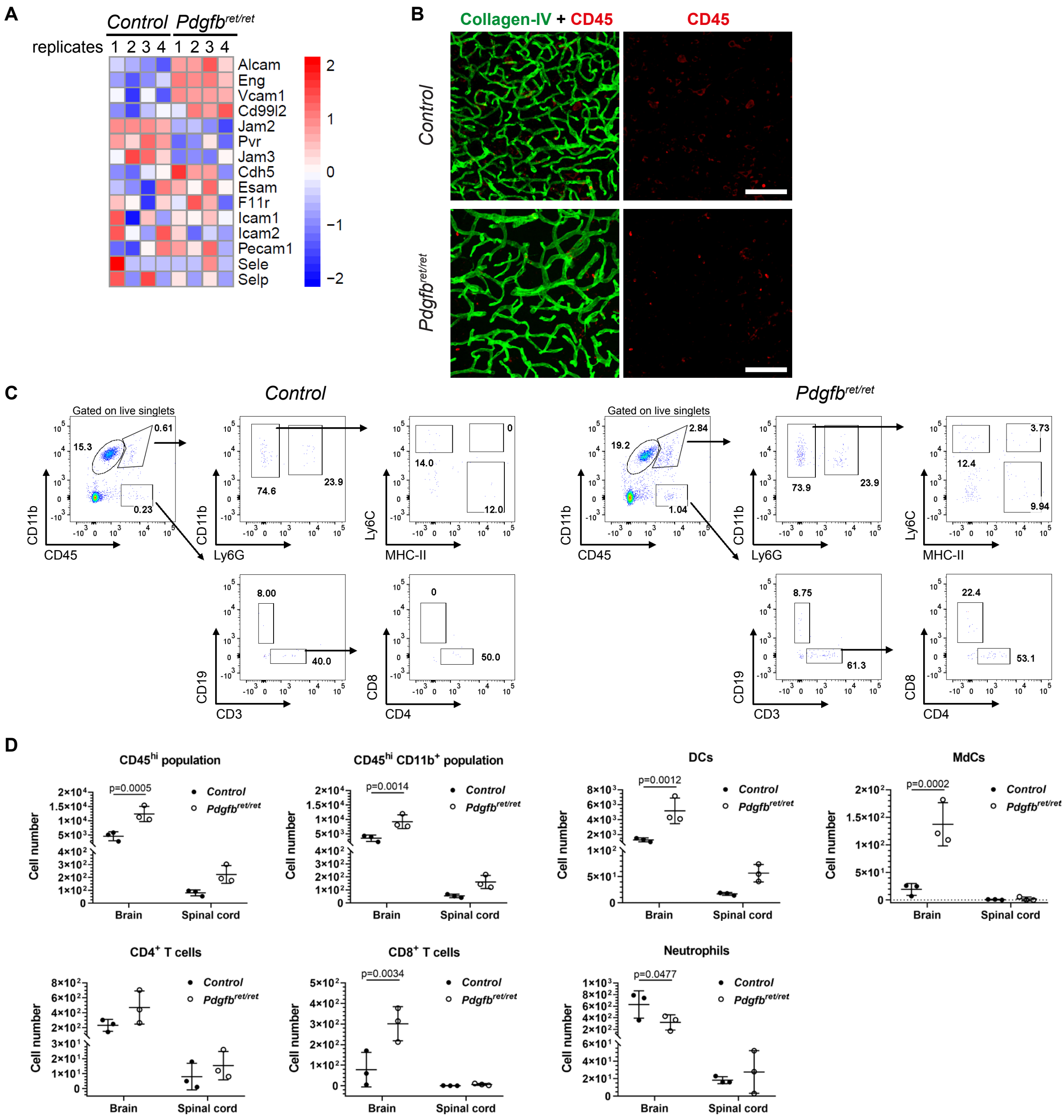
A
Control
Pdgfb ret/ret
replicates 1 2 3 4 1 2 3 4
Alcam
Eng
Vcam1
Cd99l2
Jam2
Pvr
Jam3
Cdh5
Esam
F11r
Icam1
Icam2
Pecam1
Sele
Selp
B
Collagen-IV + CD45
CD45
Control
Pdgfb ret/ret
C
Control
Pdgfb ret/ret
Gated on live singlets
CD11b
CD45
Ly6G
Ly6C
MHC-II
CD19
CD3
CD8
CD4
D
CD45hi population
CD45hi CD11b+ population
DCs
MdCs
CD4+ T cells
CD8+ T cells
Neutrophils
Cell number
Brain
Spinal cord
Control
Pdgfb ret/ret
p=0.0005
p=0.0014
p=0.0012
p=0.0002
p=0.0034
p=0.0477

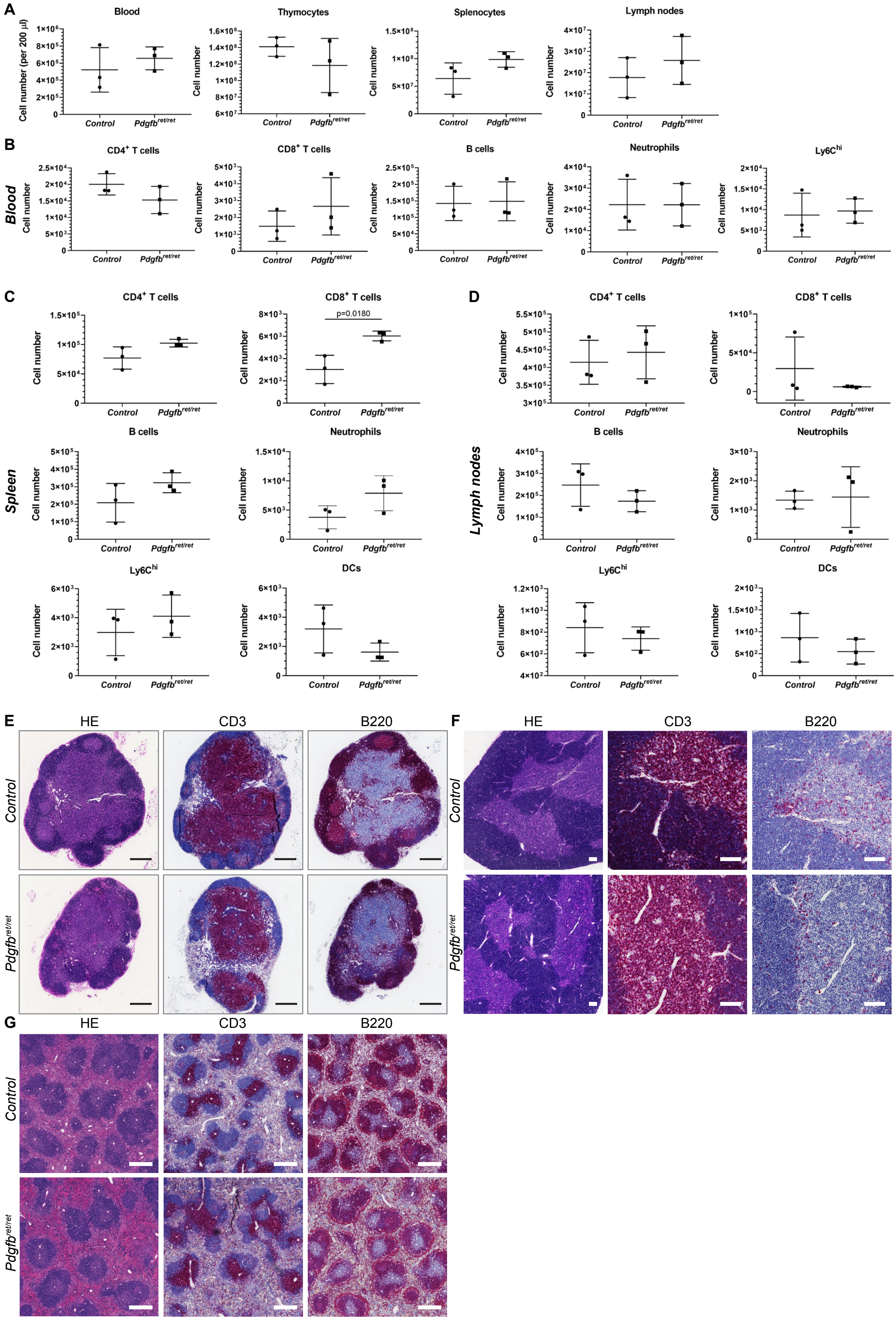
A
Blood
Thymocytes
Splenocytes
Lymph nodes
Cell number (per 200 µl)
Cell number
Control
Pdgfbret/ret
B
Blood
CD4+ T cells
CD8+ T cells
B cells
Neutrophils
Ly6Chi
C
Spleen
p=0.0180
DCs
D
Lymph nodes
E
F
G
HE
CD3
B220
Control
Pdgfbret/ret

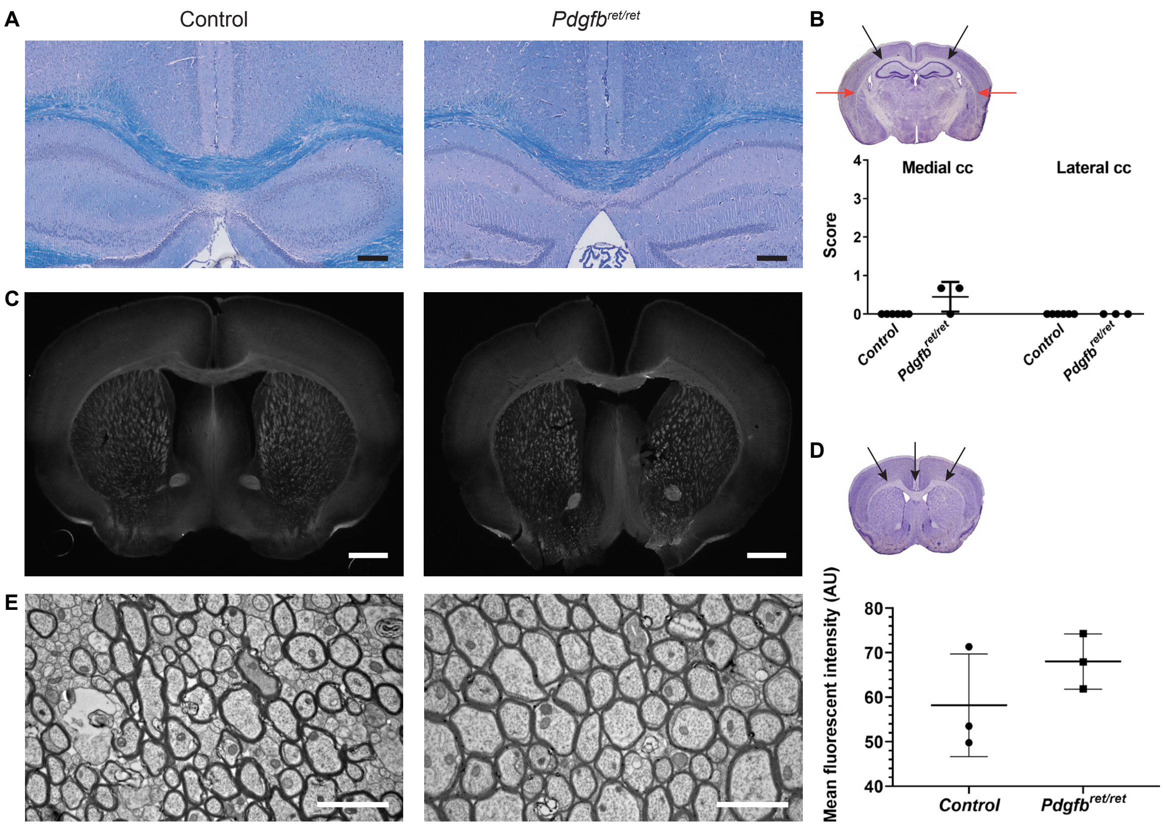
A
Control
Pdgfb ret/ret
B
Medial cc
Lateral cc
Score
4
3
2
1
0
Control
Pdgfb ret/ret
Control
Pdgfb ret/ret
C
D
E
Mean fluorescent intensity (AU)
80
70
60
50
40
Control
Pdgfb ret/ret

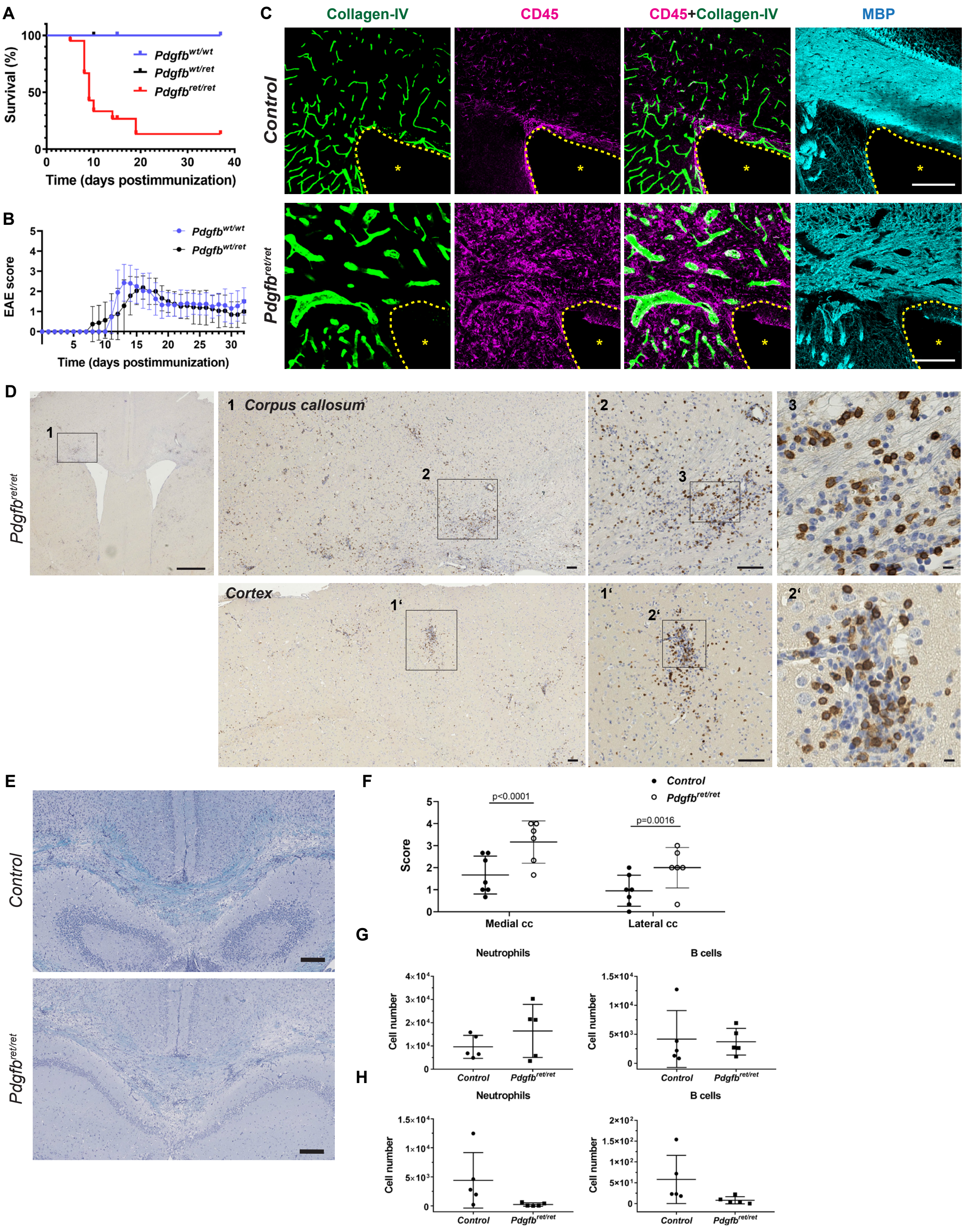
A
Survival (%)
100
50
0
0 10 20 30 40
Time (days postimmunization)
Pdgfbwt/wt
Pdgfbwt/ret
Pdgfbret/ret
B
EAE score
5 10 15 20 25 30
Time (days postimmunization)
Pdgfbwt/wt
Pdgfbwt/ret
C
Collagen-IV
CD45
CD45+Collagen-IV
MBP
Control
Pdgfbret/ret
D
Pdgfbret/ret
1 Corpus callosum
2
3
Cortex
1'
2'
E
Control
Pdgfbret/ret
F
Control
Pdgfbret/ret
Score
p<0.0001
p=0.0016
Medial cc
Lateral cc
G
Neutrophils
B cells
Cell number
Control
Pdgfbret/ret
H
Neutrophils
B cells
Cell number
Control
Pdgfbret/ret

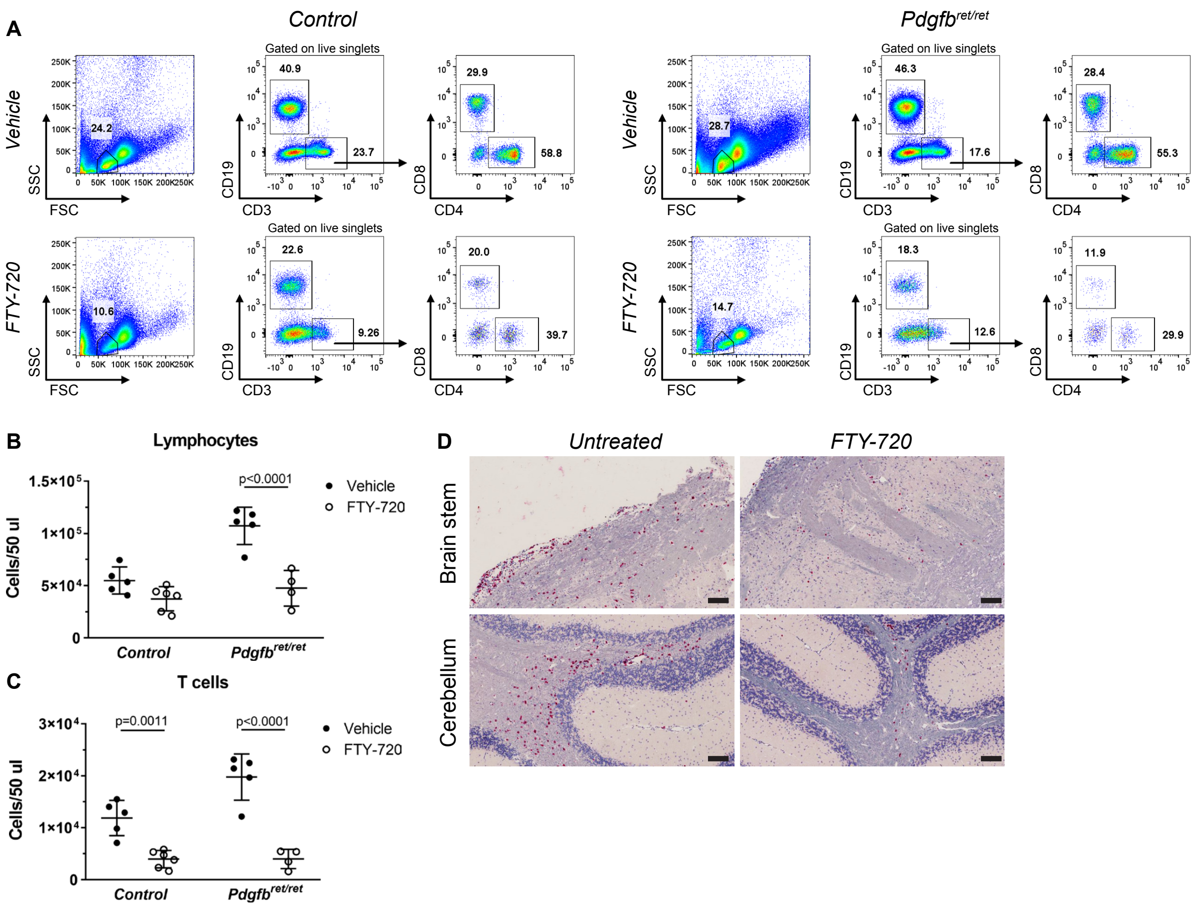
A
Control
Pdgfbret/ret
Vehicle
FTY-720
Gated on live singlets
SSC
FSC
CD19
CD3
CD8
CD4
24.2
40.9
23.7
29.9
58.8
28.7
46.3
17.6
28.4
55.3
10.6
22.6
9.26
20.0
39.7
14.7
18.3
12.6
11.9
29.9
B
Lymphocytes
Cells/50 ul
p<0.0001
Vehicle
FTY-720
Control
Pdgfbret/ret
C
T cells
p=0.0011
p<0.0001
D
Untreated
FTY-720
Brain stem
Cerebellum

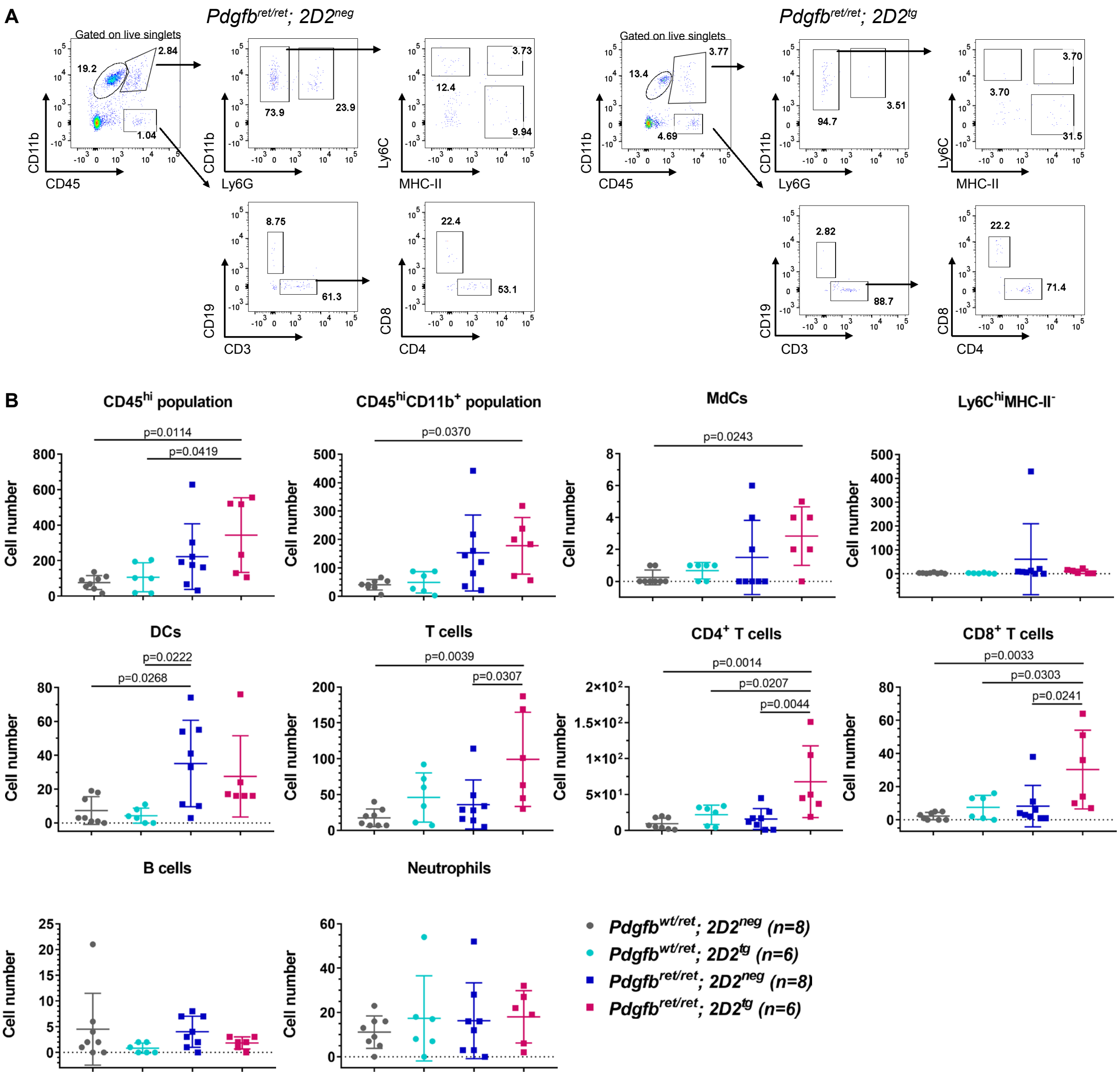
A
Pdgfbret/ret; 2D2neg
Pdgfbret/ret; 2D2tg
Gated on live singlets
CD11b
CD45
Ly6G
Ly6C
MHC-II
CD19
CD3
CD8
CD4
B
CD45hi population
CD45hiCD11b+ population
MdCs
Ly6ChiMHC-II-
DCs
T cells
CD4+ T cells
CD8+ T cells
B cells
Neutrophils
Cell number
Pdgfbwt/ret; 2D2neg (n=8)
Pdgfbwt/ret; 2D2tg (n=6)
Pdgfbret/ret; 2D2neg (n=8)
Pdgfbret/ret; 2D2tg (n=6)
